# SWELL1 channel-mediated D-2HG export promotes immune evasion and metabolic fitness in IDH-mutant glioma

**DOI:** 10.64898/2026.09.18.752798

**Authors:** Henry Yi Cheng, Lifei Ma, Jianan Chen, Jiale Huang, Li Chen, Wenrui Liu, Yongqing Liu, Runxuan Zhao, Yingzhi Ye, Wenqiang Zheng, Lisa Kratz, Maria G. Castro, Shuying Sun, Jiangjiang Zhu, Leng Han, Zhaozhu Qiu

## Abstract

Isocitrate dehydrogenase (IDH) is the most frequently mutated metabolic enzyme in human cancers. Mutant IDH produces the oncometabolite D-2-hydroxyglutarate (D-2HG), which promotes tumorigenesis in part through epigenetic alterations. Beyond its cell-autonomous effects, tumor-derived D-2HG acts as a potent immunosuppressant that establishes an immune-cold tumor microenvironment. However, the mechanism by which D-2HG is released from tumors remains largely unknown. Here, we identify the SWELL1 (LRRC8A)/LRRC8C-containing volume-regulated anion channel (VRAC) as a principal pathway for D-2HG efflux from IDH-mutant cells. Genetic deletion of SWELL1, the essential VRAC subunit, markedly reduces D-2HG release and reverses associated immunosuppression in an orthotopic mouse model of IDH-mutant glioma. Strikingly, loss of VRAC causes intracellular accumulation of D-2HG, which paradoxically limits tumor cell proliferation by driving epigenetic remodeling and mitochondrial metabolic stress. Pharmacological inhibition of VRAC with dicumarol suppresses IDH-mutant glioma growth, enhances intratumoral T cell activation, synergizes with immune checkpoint blockade, and prolongs survival in tumor-bearing mice. In human IDH-mutant lower-grade gliomas, elevated LRRC8C expression correlates with DNA hypermethylation and an immunosuppressive tumor microenvironment (TME), and predicts poor overall survival. Together, these findings establish VRAC-mediated D-2HG export as a central mechanism regulating both immune evasion and tumor cell fitness, uncovering new therapeutic opportunities across IDH-mutant cancers.

## Introduction

The most common mutation of isocitrate dehydrogenase (IDH1/2) in gliomas is a heterozygous arginine-to-histidine substitution at residue 132 of IDH1 (IDH1^R132H^), which confers a neomorphic gain-of-function that converts α-ketoglutarate (αKG) into the oncometabolite D-2-hydroxyglutarate (D-2HG) (*1*). Owing to its structural similarity to αKG, D-2HG competitively inhibits αKG-dependent enzymes, including TET family DNA hydroxylases and JmjC-domain histone demethylases, leading to widespread DNA and histone hypermethylation (*2–4*). This establishes an abnormal epigenetic landscape that leads to cell transformation (*5, 6*). In addition to its cell-autonomous effects, tumor-derived D-2HG has also emerged as a potent immunosuppressive metabolite. Accumulating to millimolar concentrations in the tumor microenvironment (TME) (*7–9*), D-2HG impairs mitochondrial respiration and effector functions of tumor-infiltrating T cells and skews tumor-associated macrophages (TAM) toward an immunosuppressive phenotype in IDH-mutant gliomas (*10–14*). Despite these profound immunosuppressive effects, how intracellular D-2HG is released into the TME remains unknown, as its negative charge precludes passive diffusion across the plasma membrane.

Through a targeted genetic screening, we identify the volume-regulated anion channel (VRAC) as a major conduit for D-2HG release from IDH-mutant cells. We demonstrate that VRAC-mediated D-2HG efflux promotes an immunosuppressive TME and unexpectedly, drives tumor cell proliferation by preventing excessive intracellular D-2HG accumulation and its anti-proliferation effects. Together, these findings establish VRAC-mediated D-2HG export as a key pleiotropic regulator of IDH-mutant glioma pathogenesis and highlight VRAC as a potential therapeutic target in IDH-mutant cancers.

### VRAC is the primary pathway for D-2HG release from mIDH tumor cells

Using human U87 glioblastoma cells harboring a heterozygous IDH1^R132H^ knock-in mutation (mIDH) as a model (*15*), we confirmed markedly elevated intracellular 2HG levels compared with isogenic wild-type (wtIDH) controls with gas chromatography coupled to mass spectrometry (GC-MS) analysis (**Fig. 1A**). Notably, after 3 days in culture, the total amount of 2HG molecules detected in the supernatants of mIDH cells exceeded intracellular levels by >25-fold (**Fig. 1B**), indicating robust 2HG release. Extracellular 2HG levels increased linearly over time, while only negligible cell death was detected by extracellular lactate dehydrogenase activity (**Fig. 1C**), suggesting that extracellular 2HG is exported by viable cells rather than released from dying cells. To characterize the mechanism of 2HG export, we subjected cell culture supernatants from U87 mIDH cells to 10-kDa MWCO filtration, which retains extracellular vesicles and proteins. 2HG passed through the filter (**Fig. 1D**), indicating that it is released as a freely soluble molecule. Cellular ATP depletion also did not alter extracellular 2HG levels (**Fig. 1E**), suggesting that 2HG export is independent of ATP-driven pumps and instead occurs via an ATP-independent transporter or channel, likely driven by the electrochemical gradient across the cell membrane. To identify this conduit, we performed an RNA interference (RNAi) screen targeting transporters (SLC22A family members, *SLC1A1*, and *SLC13A3*) previously implicated in 2HG transport (*16–19*), together with three large-pore ion channels permeable to metabolites and small signaling molecules (*20–22*) (*GJA1*, *PANX1*, and *SWELL1*). Among all candidates tested, the VRAC essential subunit SWELL1 emerged as the only major hit: its knockdown reduced extracellular 2HG levels by about half (**Fig. 1F**). VRAC regulates cell volume homeostasis during osmotic swelling by mediating chloride (Cl⁻) and organic osmolyte efflux (*20*). Beyond this canonical function, this ubiquitously expressed channel also contributes to diverse physiological and pathological processes by transporting signaling molecules, metabolites, and drugs (*23–29*). VRAC is a hetero-hexameric channel composed of SWELL1 and one or more LRRC8 paralogues (LRRC8B-E), which conder substrate selectivity(*20, 30*). We next performed RNAi knockdown of individual LRRC8 subunits to examine the VRAC composition responsible for 2HG efflux. Notably, only *LRRC8C* knockdown markedly reduced 2HG release, suggesting that SWELL1/LRRC8C-containing VRAC serves as a primary conduit for 2HG export (**Fig. 1G**). In contrast, extracellular 2HG levels modestly increased following *LRRC8D* knockdown, likely due to a compensatory increase in LRRC8C-containing VRAC upon loss of LRRC8D (*29*).

**Figure 1.**
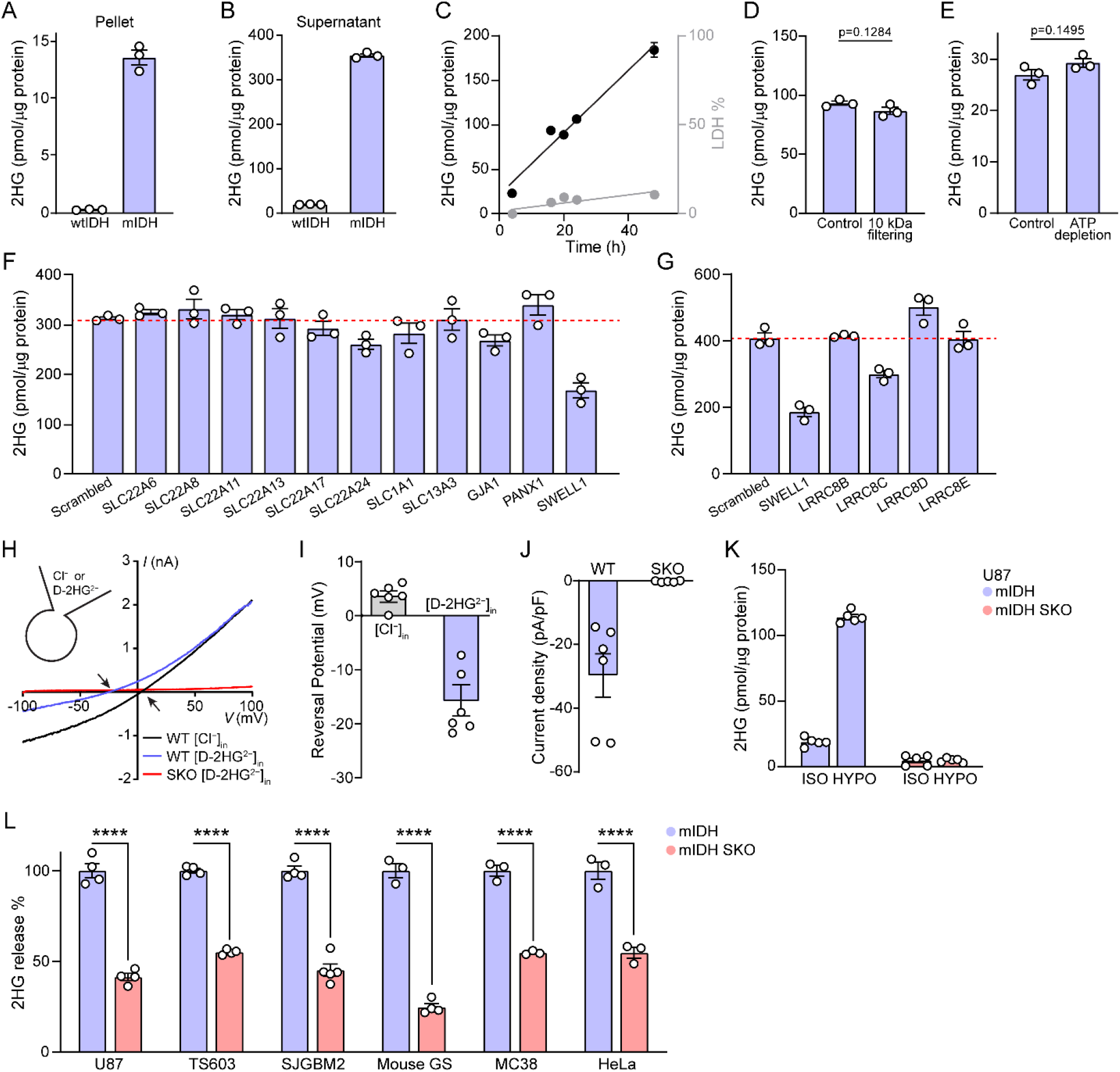
VRAC is the primary pathway for D-2HG release from mIDH tumor cells. **A–B,** 2HG concentration normalized to total protein levels in cell pellets (**A**) and culture supernatants (**B**) of U87 cells after 72 h in culture (n = 3). mIDH in (**A**) is shared with **Fig. S7A**. **C**, 2HG concentration normalized to total protein levels in cell culture supernatants of U87 cells collected at indicated times (left y axis). Corresponding supernatants were also used for LDH activity quantification (right y axis). Linear regressions were performed for the data shown (n = 3). **D,** 2HG concentration normalized to total protein levels in cell culture supernatants of U87 cells after 18 h in culture followed by 10-kDa MWCO filtering (n = 3). **E,** 2HG concentration normalized to total protein levels in cell culture supernatants of U87 cells with ATP depletion for 4 h (n = 3). Control data correspond to 4 h in Fig. 1C. **F–G,** 2HG concentration normalized to total protein levels in cell culture supernatants of siRNA-transfected U87 cells. Samples were collected 72 h after 48 h of siRNA transfections (n = 3). **H–J**, Hypotonicity-induced whole-cell electrophysiology recording in representative I-V curve (**H**), reversal potential (**I**), and current density at −100 mV (**J**) with D-2HG-based pipette solution in HeLa cells. Arrows in (**H**) indicate the reversal potentials (n = 5 to 6). **K,** 2HG concentration normalized to total protein levels in hypotonic solutions of U87 cells after 30 min of stimulation (n = 5). **L**, Relative 2HG release in cell culture supernatants of indicated cells after 48 h in culture. Data normalized to the average of mIDH samples of each cell type (n = 3 to 5). GS, gliomasphere. Data are reported as mean ± SEM. Unpaired t-test for **D**, **E**, **L**. ****p < 0.0001.

Based on its structural similarity to glutamate, a known VRAC permeant (*24*), we reasoned that D-2HG may directly permeate the VRAC pore. To test this hypothesis, we replaced intracellular Cl⁻ with equimolar D-2HG^2−^ as the sole permeant anion and activated VRAC currents by hypotonic perfusion in HeLa cells. Whole-cell patch-clamp recordings revealed small but measurable inward currents, indicating efflux of the negatively charged D-2HG from wild-type (WT) cells (**Fig. 1H**). Analysis of reversal potential shifts confirmed substantial D-2HG permeability (**Fig. 1I**), with an estimated *P*_D-2HG_/*P*_Cl_ of ∼0.16. Consistent with SWELL1 being the essential VRAC subunit, the inward currents mediated by D-2HG efflux were abolished in SWELL1 knockout (SKO) cells (**Fig. 1J**). To further validate whether VRAC mediates D-2HG release in glioma cells, we treated U87 mIDH cells with hypotonic solution for 30 min, which induced a dramatic increase in extracellular 2HG levels (**Fig. 1K**). Consistent with the VRAC dependency, this hypotonicity-induced increase in 2HG release was abolished in U87 mIDH SKO cells (**Fig. 1K** and **Fig. S1A**). Notably, extracellular 2HG levels were also reduced by ∼60% after 2 days under normal culture conditions for U87 mIDH SKO cells compared with mIDH cells (**Fig. 1L**). This result is consistent with the RNAi screen and indicates that D-2HG efflux is predominantly mediated by basal channel activity of VRAC. To test whether VRAC mediates D-2HG release more broadly, we knocked out SWELL1 using CRISPR-Cas9 in human patient-derived TS603 harboring endogenous mIDH (*31*) and pediatric glioblastoma SJGBM2 cells (*32*), and several cell lines transduced with mIDH, including human pediatric glioblastoma SJGBM2 cells (*32*), HeLa cells, primary mouse gliomasphere cells, and mouse colon adenocarcinoma MC38 cells (**Fig. S1B–D**). Deletion of SWELL1 reduced extracellular 2HG by ∼50-70% across all cell lines (**Fig. 1L**), thereby identifying VRAC as a ubiquitous and main pathway for D-2HG release from IDH-mutant cells.

### SWELL1 deletion attenuates D-2HG-driven immunosuppression in a mouse model of IDH-mutant glioma

To assess whether VRAC contributes to D-2HG release and immunosuppression *in vivo*, we implanted primary mouse mIDH gliomasphere cells orthotopically into immunocompetent C57BL/6 mice (**Fig. 2A**). Magnetic resonance imaging (MRI) analysis at 28 days post-implantation revealed markedly smaller tumors in mice bearing SWELL1 knockout (SKO) gliomas compared with mIDH controls (**Fig. 2B** and **Fig. S2A**). Consistent with reduced tumor burden, mice harboring mIDH SKO tumors exhibited prolonged survival, with a median survival (MS) of 45 days versus 34 days in control mice (**Fig. 2C**). D-2HG levels in tumor-adjacent brain tissues collected at endpoint were substantially reduced in SKO tumors (**Fig. 2D**), supporting a role for VRAC in D-2HG release into the TME. To characterize the immune profile within the TME, we first performed flow cytometry (FACS) analysis of tumor-infiltrating myeloid cells (**Fig. S2B** and **S2C**). Compared with mIDH controls, SKO tumors contained reduced tumor-associated neutrophils (TANs) and monocytic myeloid-derived suppressor cells (m-MDSCs) populations (**Fig. S2D**). Notably, microglia and TAMs within mIDH SKO tumors exhibited an enhanced activation phenotype, as indicated by decreased expression of Arginase-1 and PD-L1 and increased populations expressing high surface levels of antigen-presenting molecules CD86 and I-A/I-E (CD86^hi^ and I-A/I-E^hi^) (**Fig. 2E–H** and **Fig. S2E–F**).

**Figure 2.**
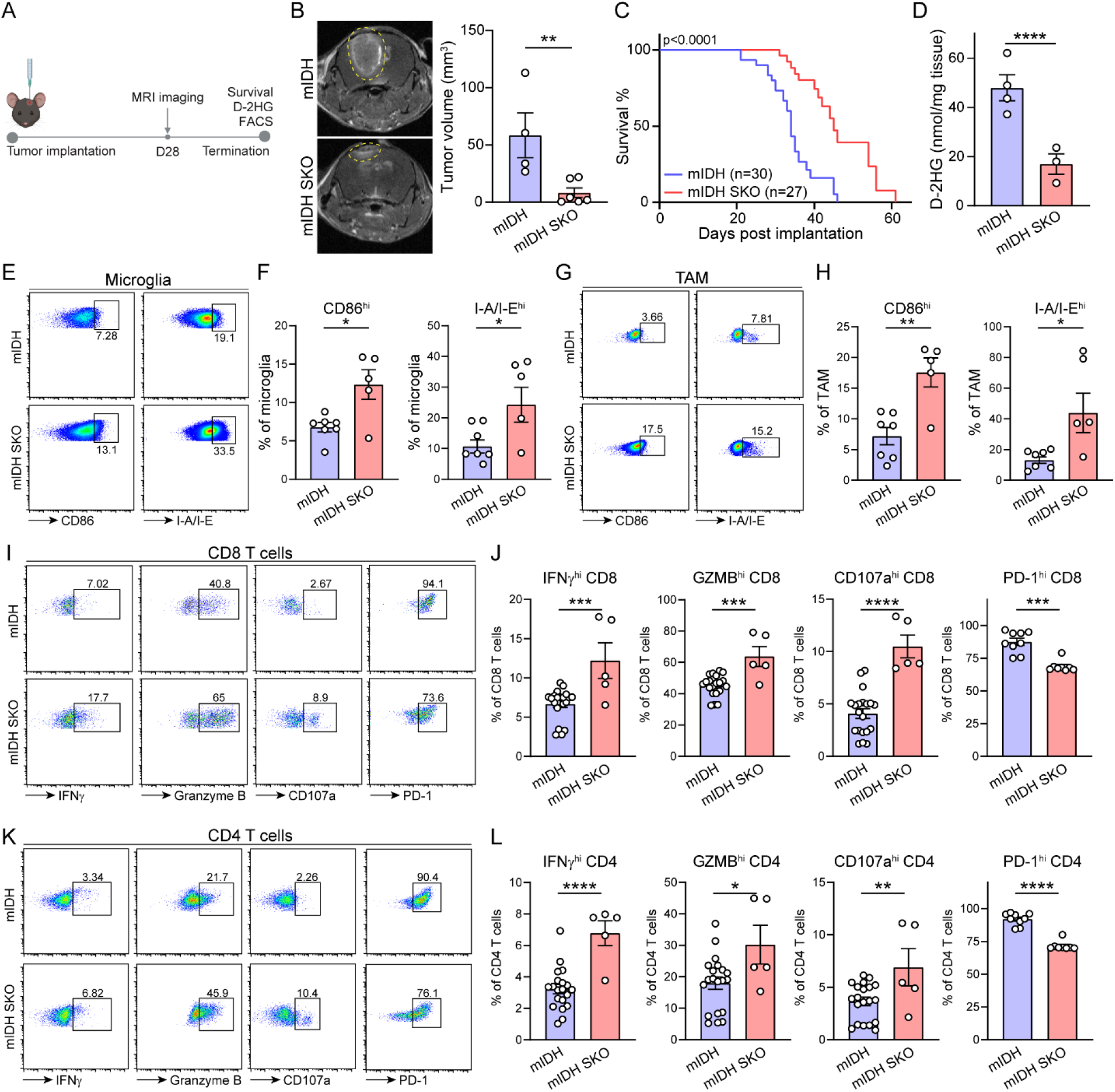
SWELL1 deletion attenuates D-2HG-driven immunosuppression in a mouse model of IDH-mutant glioma. **A**, Workflow summary for the syngeneic orthotopic mouse glioma model. **B**, Representative MRI images and tumor volume quantification of mouse gliomasphere tumors on day 28 (n = 4 to 6). **C**, Survival of mice implanted with 5 x 10^4^ mouse gliomasphere cells. **D**, D-2HG concentration normalized to tissue weight in tumor-adjacent brain tissues from mice harboring mouse gliomasphere tumors (n = 3 to 4). **E–H**, FACS analysis (left) and quantification (right) of tumoral microglia (**E–F**) or tumor-associated macrophages (TAMs) (**G–H**) for high surface CD86 and I-A/I-E populations (n = 5 to 8). **I–L**, FACS analysis (left) and quantification (right) of tumoral CD8+ T cells (**I–J**) or CD4+ T cells (**K–L**) for high IFNγ, Granzyme B, surface CD107a, and PD-1 populations (n = 5 to 20). GZMB, granzyme B. Data are reported as mean ± SEM. Unpaired t-test for **B, D, F, H, J, L.** Mantel-Cox test for **C**. *p < 0.05, **p < 0.01, ***p < 0.001, and ****p < 0.0001.

Tumor-associated microglia and macrophages are key antigen-presenting cells that drive tumoral lymphocyte activation (*10*). We therefore profiled CD8^+^ and CD4^+^ T cell populations within the TME (**Fig. S2G**). Consistent with enhanced myeloid activation, mIDH SKO tumors exhibited increased CD8⁺ T cell infiltration accompanied by a reduction in regulatory T cells (Tregs) (**Fig. S2H**). Moreover, these tumors displayed a marked shift toward a functionally activated T cell phenotype, characterized by increased frequencies of IFNγ^hi^, granzyme B^hi^, surface CD107a^hi^ CD8^+^ and CD4^+^ T cells, together with reduced PD-1^hi^ populations and elevated expression of the effector cytokines IL-2 and TNFα (**Fig. 2I–L** and **Fig. S2I–J**), indicating enhanced cytotoxic and effector T cell responses. To determine whether these enhanced immune responses resulted from reduced D-2HG release or increased intrinsic immunogenicity of tumor cells, we performed RNA sequencing (RNA-seq) on primary mouse gliomasphere cells and grouped differentially expressed genes (DEGs) into inflammation, MHC antigen presentation, and immune checkpoint ligand modules. Importantly, mIDH SKO cells showed no systematic alterations across these programs compared to mIDH cells, with only modest increases in *Sting* and *Casp1* expression and a modest decrease in *Ccl5* (**Fig. S2K**). Together, these findings demonstrate that SWELL1-dependent D-2HG efflux suppresses anti-tumor immunity and that its loss reprograms the TME toward enhanced immune activation in IDH-mutant gliomas.

### Low SWELL1 expression correlates with an immunoactive TME in human IDH-mutant gliomas

To further examine the role of VRAC in the TME, we analyzed single-cell RNA-sequencing data from 12 IDH-mutant glioma patients (*33*). Dozens of distinct cell populations were identified spanning malignant, stromal, and immune lineages (**Fig. S3A** and **S3B**). Notably, *LRRC8C* transcripts were rarely detected in malignant cells, likely due to limited sequencing depth. In contrast, *SWELL1* was more robustly expressed, allowing reliable patient stratification. Accordingly, tumors were divided into *SWELL1*-high and *SWELL1*-low groups based on the median *SWELL1* expression level in malignant cells. Compared with *SWELL1*-high tumors, *SWELL1*-low tumors contained increased proportions of microglia and macrophages (**Fig. S3C**), accompanied by upregulation of genes involved in immune-response pathways, including defense response, inflammation, and immune cell differentiation and activation (**Fig. S3D** and **S3E**). In addition, malignant cells and cycling cells, a small but functionally important population that drives tumor proliferation (*34*), also exhibited enrichment of inflammatory gene signatures, including cytokine activity and signaling. Consistent with our findings in the syngeneic mouse glioma model (**Fig. 2**), these results indicate that low *SWELL1* expression is associated with an immunoactive TME in human IDH-mutant gliomas. Concordantly, macrophages within *SWELL1*-high tumors were enriched for TGFβ-responsive gene programs, which have been linked to D-2HG-induced immunosuppressive phenotype (*10*). Microglia in *SWELL1*-high tumors were enriched for nutrient-and hormone-response pathways, the roles of which in IDH-mutant gliomas remain unclear. Notably, *SWELL1*-high tumors contained higher proportions of malignant and cycling cells (**Fig. S3C**). These malignant cells exhibited enriched cell proliferation and TGFβ signaling pathways (**Fig. S3F**), features often associated with more aggressive tumor behavior (*35*), while cycling cells were enriched for multiple mitotic pathways (**Fig. S3G**). These results suggest that VRAC may contribute to mIDH tumor cell proliferation while simultaneously shaping the TME.

### VRAC-mediated D-2HG efflux limits intracellular D-2HG levels and promotes cell proliferation by epigenetic remodeling in mIDH glioma cells

To investigate a potential role for VRAC in mIDH glioma cell proliferation, we evaluated growth activity of primary mouse gliomasphere cells by Ki67 staining and colony-formation assays. Consistent with previous reports (*36–39*), mIDH gliomasphere cells exhibited lower proliferative capacity than wtIDH cells (**Fig. 3A** and **3B**), supporting a complex role for D-2HG in glioma biology—promoting oncogenesis during tumor initiation while constraining cell growth during tumor progression (*5, 6, 32, 36–38, 40*). Deletion of SWELL1 further reduced proliferation of mIDH cells (**Fig. 3A** and **3B**), indicating a role for VRAC in promoting cell proliferation. Notably, this effect was specific to the mIDH background, as SWELL1 loss did not impair growth in wtIDH cells (**Fig. S4A**). Given the lack of efficient enzymatic degradation (*41*), we hypothesized that VRAC-mediated efflux is a major pathway for D-2HG clearance and that its loss leads to intracellular accumulation and growth suppression. To test this hypothesis, we measured intracellular 2HG levels and observed a ∼2-fold increase in mIDH SKO cells compared to controls (**Fig. 3C**), consistent with impaired D-2HG export. Similar results were obtained using an independent CRISPR gRNA targeting SWELL1 (mIDH SKO2) (**Fig. S4B**). These findings support a model in which VRAC-mediated D-2HG release relieves intracellular D-2HG–dependent growth suppression, thereby promoting mIDH cell proliferation.

**Figure 3.**
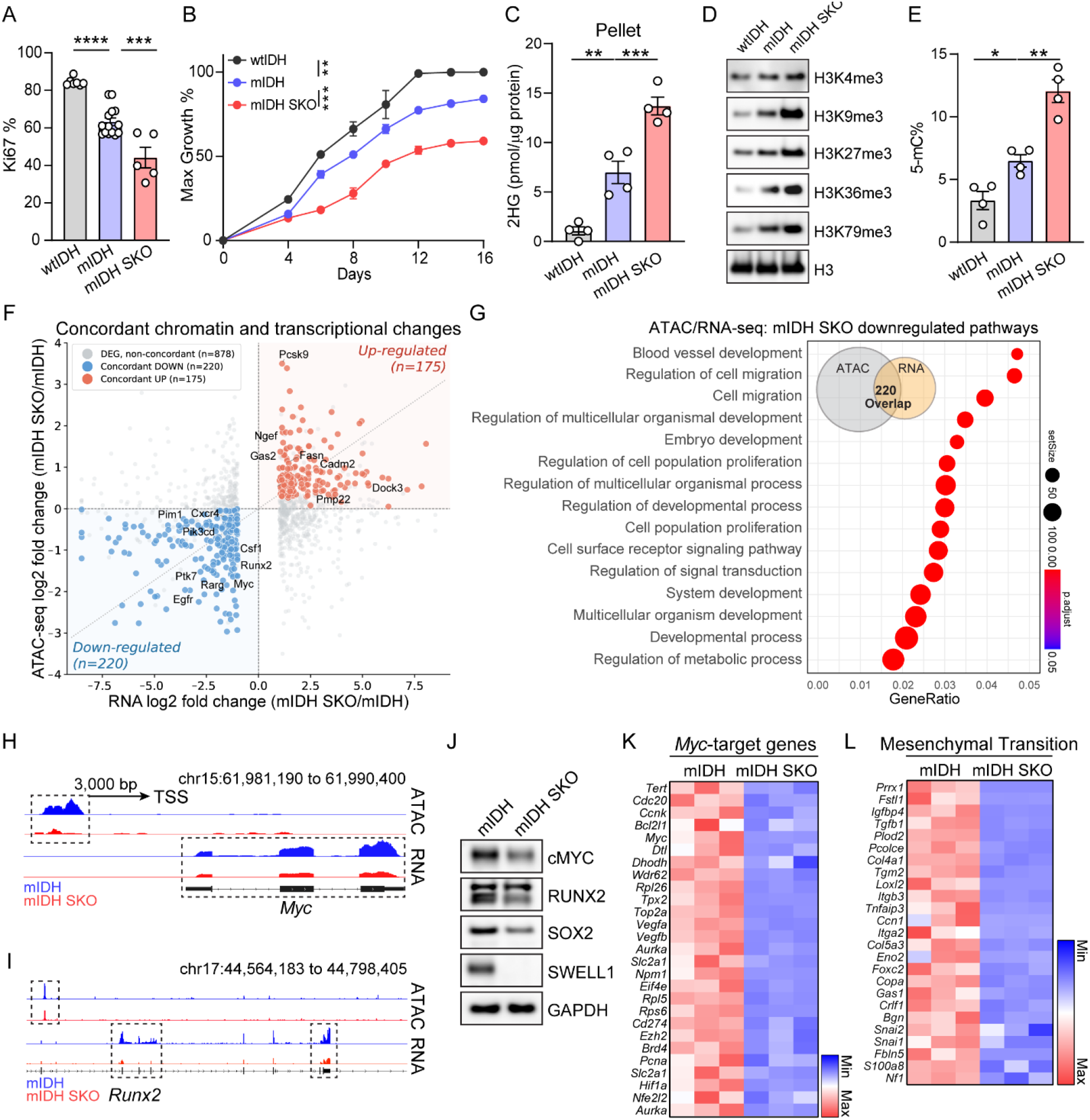
VRAC-mediated D-2HG efflux limits intracellular D-2HG accumulation and promotes cell proliferation by shaping D-2HG-driven epigenetic remodeling in mIDH glioma cells. **A**, FACS quantification of Ki67 positivity of mouse gliomasphere cells (n = 5 to 14). **B**, Colony formation assay for mouse gliomasphere cells (n = 3). Data normalized to the maximum absorbance within the comparison. wtIDH data is shared with **Fig. S4A**. **C**, 2HG concentration normalized to total protein levels in mouse gliomasphere cell pellets (n = 4). wtIDH and mIDH are shared with **Fig. S4B**. **D**, Immunoblotting of isolated histones from mouse gliomasphere cells for H3 trimethylations (n = 3). H3 is shared with **Fig. S4C**. **E**, 5-methylcytosine (5-mC) quantification normalized to total cytosine in mouse gliomasphere cells (n = 4). **F**, Scatter plot of concordant chromatin accessibility (ATAC-seq) and transcriptional (RNA-seq) changes in mIDH SKO vs. mIDH cells, identifying 220 concordantly downregulated and 175 concordantly upregulated gene loci, with select oncogenic (e.g., *Myc*, *Egfr*, *Runx2*) and developmental (e.g., *Pcsk9*, *Shank2*) genes labeled. **G**, Selected gene ontology biological process terms enriched among the 220 concordantly downregulated genes. Inset Venn diagram shows the 220-gene overlap between ATAC-seq (n = 4) and RNA-seq datasets (n = 3). **H–I**, IGV profiles of representative ATAC-seq and RNA-seq signals for *Myc* (**H**) or *Runx2* (**I**) in mouse gliomasphere cells (ATAC-seq: n = 4; RNA-seq: n = 3). Loci are oriented such that the TSS is shown on the left. **J**, Immunoblotting of mouse gliomasphere cells for cMYC, RUNX2, SOX2, SWELL1, and GAPDH in mIDH and mIDH SKO cells (n = 3). **K**, Heatmaps of MYC-target gene expression within mouse gliomasphere RNA-seq (n = 3). Data presented in z-scores calculated from FPKM. **L**, Heatmaps of mesenchymal transition gene expression within mouse gliomasphere RNA-seq (n = 3). Data presented in z-scores calculated from FPKM. Data are reported as mean ± SEM. One-way ANOVA with Sidak’s test for **A, C, E**. Two-way ANOVA with Sidak’s test for **B.** *p < 0.05, **p < 0.01, ***p < 0.001, and ****p < 0.0001.

To investigate the underlying mechanisms, we first assessed epigenetic changes, as D-2HG competitively inhibits αKG-dependent demethylases and promotes hypermethylation of histones and DNA (*2–4*). As expected, mIDH gliomasphere cells exhibited globally elevated histone H3 trimethylation at lysine residues K4, K9, K27, K36, and K79 compared with wtIDH cells (**Fig. 3D**). In contrast, histone H3 acetylation at K9 and K14 remained unchanged (**Fig. S4C**). Consistent with elevated intracellular D-2HG levels, the trimethylation marks were further increased in mIDH SKO cells relative to mIDH controls (**Fig. 3D**). Importantly, histone hypermethylation was specific to the mIDH context, as SWELL1 deletion in wtIDH cells did not alter H3 trimethylation (**Fig. S4D**). Furthermore, total DNA methylation, measured by 5-methylcytosine (5-mC), was also elevated upon SWELL1 deletion in mIDH cells (**Fig. 3E**).

To investigate the transcriptional consequences of epigenetic remodeling, we analyzed the RNA-seq data from wtIDH, mIDH, and mIDH SKO gliomasphere cells. Consistent with previous reports (*32, 36–38*), pathway enrichment analysis revealed upregulation of DNA repair pathways as well as downregulation of cell proliferation and growth factor signaling pathways in mIDH cells relative to wtIDH cells (**Fig. S4E–G**). We next focused on DEGs common to both the mIDH SKO versus mIDH and the mIDH versus wtIDH comparisons (**Fig. S4H**). These genes were enriched for neuronal differentiation programs and depleted for pathways related to cell proliferation, migration, and metabolism (**Fig. S4I** and **S4J**), suggesting that loss of SWELL1 promotes cellular differentiation and reduces proliferative capacity in mIDH cells.

DNA hypermethylation and repressive histone methylation marks (e.g., H3K9me3 and H3K27me3) are commonly associated with transcriptional silencing, whereas activating histone methylation marks (e.g., H3K4me3 and H3K36me3) can promote transcription (*42*). To link transcriptional changes with epigenetic alterations, we performed assay for transposase-accessible chromatin-sequencing (ATAC-seq) to assess chromatin accessibility and focused on differentially accessible regions associated with protein-coding genes in mIDH SKO gliomasphere cells versus mIDH controls. By overlapping these genomic loci with DEGs identified by RNA-seq, we found 220 genes exhibiting concordant reductions in chromatin accessibility and transcriptional output in mIDH SKO cells (**Fig. 3F**). These genes were enriched in pathways related to cell proliferation, migration, and signal transduction (**Fig. 3G**), including several oncogenes critical for glioma pathogenesis (*39, 43–45*): *Cd34*, *Ptk7*, *Peg10*, *Myc*, and *Runx2* (**Fig. 3H** and **3I** and **Fig. S5A–C**). Among these, *Myc* and *Runx2* encode key transcription factors that drive tumor progression and mesenchymal transition (*39, 44*). Reduced protein expression of MYC and RUNX2 (the lower molecular weight isoform 1) (*46*) in mIDH SKO cells was confirmed by immunoblotting (**Fig. 3J**). This was accompanied by concordant downregulation of MYC target genes and mesenchymal transition programs in the RNA-seq data (**Fig. 3K** and **3L**). In parallel, 175 genes exhibited concordant increases in chromatin accessibility and transcript abundance in mIDH SKO cells and were enriched for neuronal differentiation and function (**Fig. 3F** and **Fig. S5D**). Consistently, RNA-seq data revealed a broader spectrum of upregulation in established regulators of neuronal differentiation (*6, 47–50*) such as *Mef2c, Egr3, Nefl, Ngef, Dlx1,* and *Dlx2* (**Fig. S5E**). Among these, EGR3 and DLX1/DLX2 are transcription factors whose regulon activity was significantly elevated in mIDH SKO cells (**Fig. S5F**), indicating activation of neuronal differentiation programs. Glioma stemness and differentiation program are largely orchestrated by the stemness-associated transcriptional factor SOX2 (*51*). We observed that SOX2 protein expression was reduced in mIDH SKO cells, accompanied by concordant downregulation of its transcriptional targets (**Fig. 3J** and **Fig. S5G**). Together, these findings suggest that loss of SWELL1 in mIDH glioma cells shifts transcriptional programs away from stemness and proliferation toward neuronal differentiation, consistent with enhanced intracellular D-2HG-driven epigenetic reprogramming.

### VRAC-mediated D-2HG release alleviates mitochondrial metabolic stress in mIDH cells

In addition to its role in epigenetic remodeling, D-2HG induces mitochondrial metabolic reprogramming, in part by depleting αKG and other intermediates of the tricarboxylic acid (TCA) cycle (*52–56*). Accordingly, targeted metabolomic profiling using combined GC-MS and liquid chromatography (LC) coupled to MS (LC-MS) revealed marked reductions in multiple TCA cycle intermediates, including citrate, αKG, succinate, and fumarate, in mIDH gliomasphere cells compared to wtIDH controls (**Fig. 4A**), whereas glycolytic and purine catabolism intermediates were largely unchanged (**Fig. S6A** and **S6B**). Consistent with increased intracellular D-2HG accumulation, these TCA cycle intermediates were further decreased in mIDH SKO cells (**Fig. 4A** and **Fig. S6C**). Because TCA cycle integrity is essential for mitochondrial respiration, we next assessed mitochondrial function using Seahorse analysis. In line with the metabolomic data, mIDH cells exhibited reduced basal respiration, ATP production, and maximal respiratory capacity compared with wtIDH controls, with mIDH SKO cells showing an even greater reduction (**Fig. 4B** and **4C** and **Fig. S6D–S6E**). These results indicate that VRAC-mediated D-2HG efflux mitigates D-2HG-driven metabolic stress.

**Figure 4.**
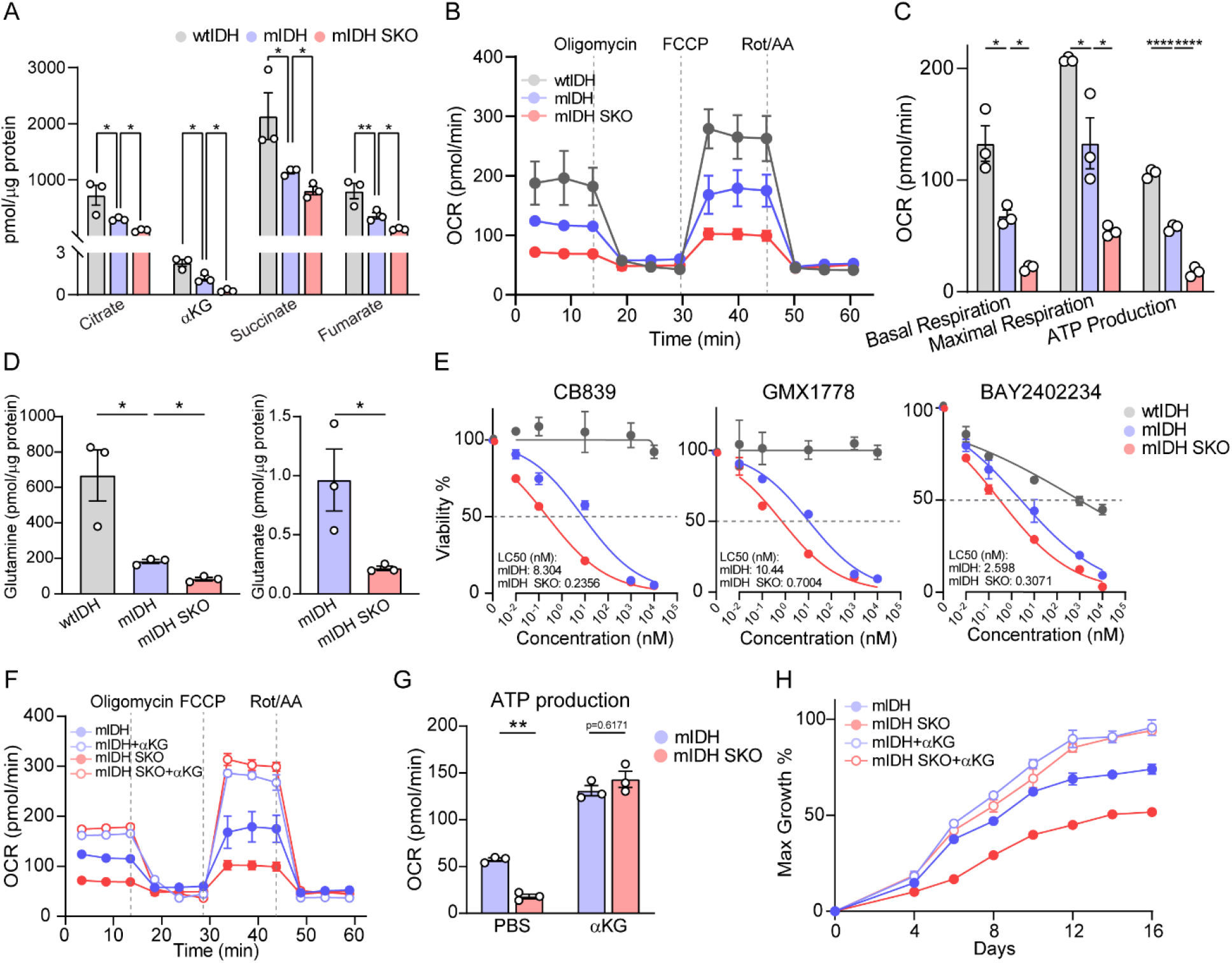
VRAC-mediated D-2HG efflux mitigates mitochondrial metabolic stress in mIDH glioma cells. **A**, TCA cycle metabolite concentrations normalized to total protein levels in mouse gliomasphere cell pellets (n = 3). mIDH and mIDH SKO are shared with **Fig. S6C**. **B–C**, Seahorse analysis of cellular oxygen consumption rate (OCR) kinetics. Representative OCR trace (**B**) and quantifications (**C**) measured in mouse gliomasphere cells (n = 3). mIDH is shared with **Fig. S6D–E**. mIDH and mIDH SKO are shared with Fig. 4F**–G** **and S6G**. **D,** Glutamine and glutamate concentrations normalized to total protein levels in mouse gliomasphere cell pellets (n = 3). mIDH is shared with **Fig. S6F**. **E**, LC50 assays for mouse gliomasphere cells after treatments of indicated compound concentrations for 72 h (n = 3). Non-linear regressions with normalized response were performed for the data shown. LC50 concentrations were determined as concentration at 50% viability. **F–G**, Seahorse analysis of cellular oxygen consumption rate (OCR) kinetics. Representative OCR trace (**F**) and ATP production (**G**) measured in mouse gliomasphere cells supplemented with 10 mM αKG (n = 3). Untreated mIDH and mIDH SKO are shared with Fig. 4B**–C**. **H**, Colony formation assay for mouse gliomasphere cells supplemented with 10 mM αKG (n = 3). Data normalized to the maximum absorbance within the comparison. Data are reported as mean ± SEM. Unpaired t-test for **A, D.** One-way ANOVA with Sidak’s test within each parameter for **C**. Two-way ANOVA with Sidak’s test for **G**. *p < 0.05, **p < 0.01, and ***p < 0.001.

When TCA cycle intermediates are depleted, glutaminolysis serves as a major anaplerotic pathway to replenish αKG (*52–54*). Consistently, intracellular glutamine levels were reduced in mIDH cells compared to wtIDH controls, with even greater reductions in both glutamine and glutamate observed in mIDH SKO cells relative to mIDH cells (**Fig. 4D** and **Fig. S6F**). These findings suggest an increased reliance on compensatory glutaminolytic metabolism to sustain energy production upon intracellular D-2HG accumulation. Increased metabolic stress in IDH-mutant gliomas creates dependence on pathways such as glutaminolysis, NAD⁺ salvage, and pyrimidine synthesis to sustain growth, thereby generating therapeutically exploitable vulnerabilities (*52–54, 57, 58*). We therefore performed median lethal concentration (LC_50_) analyses using inhibitors targeting these pathways (**Fig. 4E**). While wtIDH cells were largely insensitive to the glutaminolysis inhibitor CB839 and the NAD^+^ salvage inhibitors GMX1778 and exhibited only modest sensitivity to the pyrimidine synthesis inhibitor BAY2402234 (LC_50_ ∼1 μM), mIDH cells displayed markedly increased sensitivity to all three compounds, with LC_50_ values ranging from 2-10 nM. Notably, SWELL1 deletion further sensitized mIDH cells, resulting in sub-nanomolar LC_50_ values, consistent with heightened dependence on compensatory metabolic pathways.

Because αKG depletion has been implicated in driving metabolic stress in IDH-mutant gliomas (*53–56*), we next tested whether αKG supplementation could rescue mitochondrial dysfunction. Supplementation of culture media with αKG markedly improved basal respiration, ATP production, and maximal respiratory capacity in both mIDH and mIDH SKO gliomasphere cells (**Fig. 4F** and **4G** and **Fig. S6G–I**). In parallel, αKG supplementation increased proliferation and restored growth to comparable levels, consistent with the reversal of both metabolic and epigenetic alterations (*2, 3*) (**Fig. 4H**). Together, these findings demonstrate that VRAC-mediated D-2HG efflux mitigates mitochondrial metabolic stress and associated metabolic vulnerabilities in mIDH glioma cells.

### SWELL1 deletion suppresses tumor growth and extends survival in a patient-derived IDH-mutant glioma xenograft mouse model

We next extended our findings from primary mouse gliomasphere cells to human glioma cells harboring mIDH, including U87, SJGBM2, and TS603 cells. Consistent with the reduced export (**Fig. 1K**), SWELL1 deletion in all three cell types led to increased intracellular D-2HG levels (**Fig. 5A** and **Fig. S7A**), accompanied by elevated histone H3 trimethylation (**Fig. 5B** and **Fig. S7B**). In TS603 SKO cells, this was associated with reduced MYC and SOX2 expression (**Fig. 5C**), concordant downregulation of their target genes, and upregulation of differentiation-associated gene programs (**Fig. 5D–F**), recapitulating the transcriptional changes observed in mouse mIDH SKO cells (**Fig. 3** and **Figs. S4–5**). Accordingly, loss of SWELL1 impaired growth of human mIDH glioma cells, as evidenced by reduced Ki67 positivity (**Fig. 5G** and **Fig. S7C**), and increased sensitivity of TS603 cells to CB839, GMX1778, and BAY2402234 (**Fig. 5H**). Finally, we performed orthotopic xenograft experiments by intracranially implanting human patient-derived TS603 cells into immunodeficient *SCID* mice. MRI analysis at day 56 post-implantation revealed markedly reduced tumor growth in mice bearing TS603 SKO cells compared to controls (**Fig. 5I** and **Fig. S7D**). Consistently, these mice exhibited significantly prolonged survival, with a median survival of 99 days versus 76 days in controls (**Fig. 5J**). Together, these results demonstrate that, as observed in mouse gliomasphere cells, VRAC-mediated D-2HG export limits intracellular D-2HG accumulation and thereby regulates epigenetic reprogramming, metabolic vulnerabilities, and proliferation in human mIDH glioma cells, as well as tumor growth in a patient-derived xenograft model.

**Figure 5.**
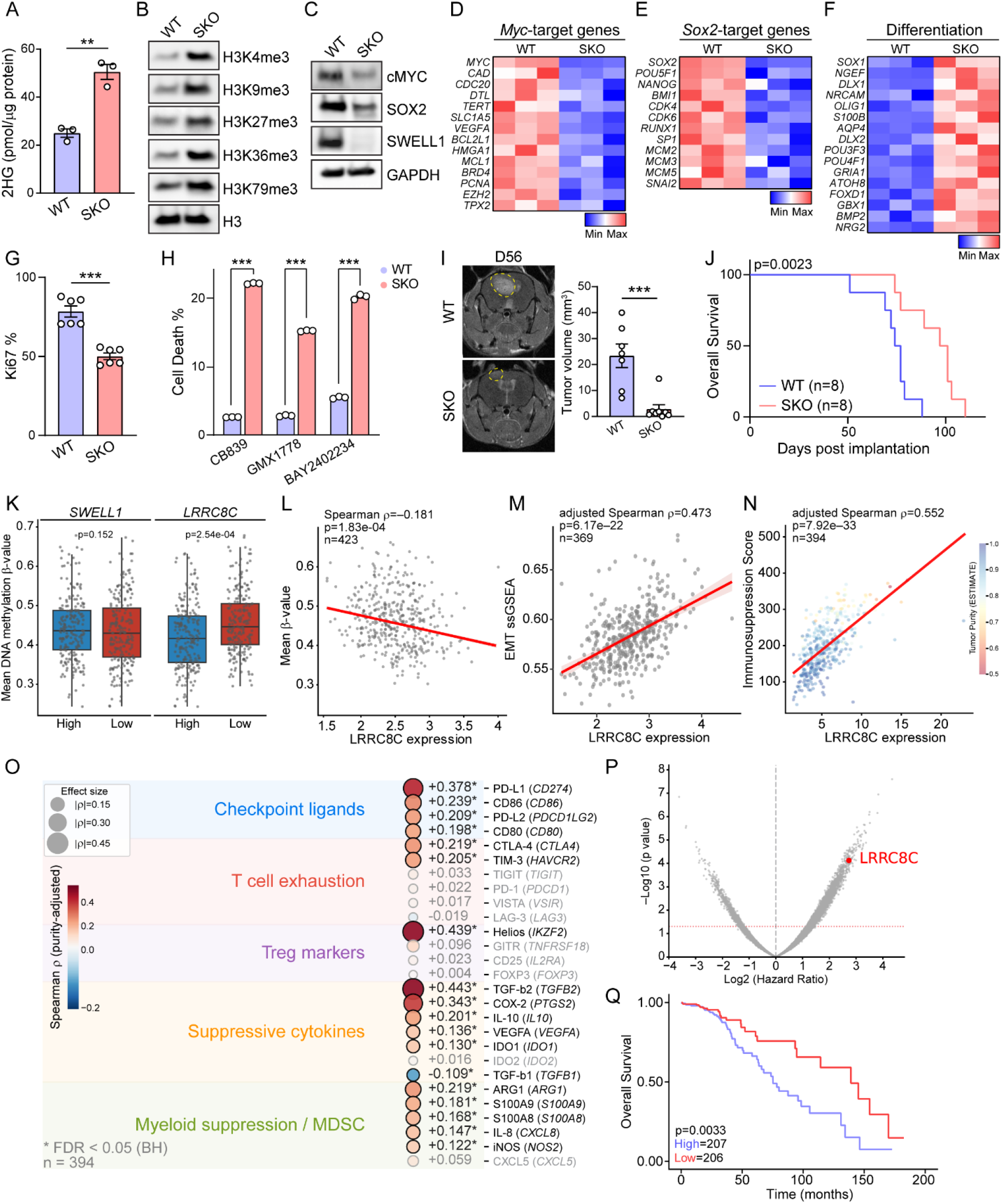
SWELL1 deletion suppresses tumor growth and extends survival in a patient-derived IDH-mutant glioma model, and high *LRRC8C* expression marks hypomethylated, immunosuppressive, poor-prognosis human tumors. **A**, 2HG concentration normalized to total protein levels in TS603 cell pellets (n = 3). **B–C**, Immunoblotting of TS603 isolated histones for H3 trimethylations (**B**) or whole cell lysates for cMYC and SOX2 expressions (**C**) (n = 3). SWELL1 and GAPDH in (**C**) are shared with **Fig. S1A**. **D–F**, Heatmaps of TS603 MYC-(**D**), SOX2-target (**E**), or cellular differentiation (**F**) gene expressions assayed by qRT-PCR (n = 3). Data presented in z-scores. **G**, FACS quantification of Ki67 positivity of TS603 cells (n = 6). **H**, FACS quantification of Live/Dead Aqua positivity of TS603 cells treated with CB839 (2 nM), GMX1778 (0.5 nM), or BAY2402234 (0.04 nM) for 72 h (n = 3). **I**, Representative MRI images and tumor volume quantification of TS603 tumors at day 56 (n = 7 to 8). **J**, Survival of *SCID* mice implanted with 1 x 10^5^ TS603 cells. **K**, Box plots comparing mean DNA methylation β-values at mIDH-sensitive 1308 CpG sites between *SWELL1*-high vs. -low and *LRRC8C*-high vs. -low IDH-mutant LGG samples from TCGA-LGG. **L–M**, Scatter plot showing the correlation between *LRRC8C* expression (Log2(TPM+1)) and mean DNA methylation β-value (**L**) and grade-adjusted epithelial-to-mesenchymal transition (EMT) ssGSEA score (**M**) in IDH-mutant LGG samples from TCGA-LGG. **N**, Scatter plot showing a significant positive correlation between *LRRC8C* expression (Log2(TPM+1)) and immunosuppression score (derived from ranked expression of *TGFB2, IL10, IDO1, ARG1, CD274, IKZF2, HAVCR2*, and *CTLA4*) in IDH-mutant LGG samples from TCGA-LGG, with tumor purity indicated by color gradient. **O**, Bubble plot of purity-adjusted Spearman correlations between *LRRC8C* expression (Log2(TPM+1)) and immunosuppressive gene signatures in IDH-mutant LGG, including checkpoint ligands, T cell exhaustion markers, Treg markers, suppressive cytokines, and myeloid suppression/MDSC markers. Genes without statistical significance are colored in light gray. **P**, Volcano plot of genome-wide multivariate Cox regression using whole transcriptome genes as prognostic factors in IDH-mutant LGG samples from TCGA-LGG. Gray and red dashed line denotes Log2HR = 0 and statistical significance threshold respectively. **Q**, Kaplan-Meier survival curves for IDH-mutant LGG patients stratified by median *LRRC8C* expression from TCGA-LGG. Data are reported as mean ± SEM. Unpaired t-test for **A, G, H, I.** Log-rank (Mantel-Cox) test for **J**, **Q**. Welch’s t-test for **K**. Spearman rank correlation for **L**, **M**. Spearman rank correlation with purity-adjusted partial Spearman correlation for **N**. purity-adjusted partial Spearman correlation with Benjamini-Hochberg FDR correction for **O**. Wald test for **P**. **p < 0.01, ***p < 0.001, and ****p < 0.0001.

### High LRRC8C expression associates with DNA hypomethylation, immunosuppression, and poor survival in IDH-mutant gliomas

To assess the clinical relevance of our findings, we interrogated the TCGA lower-grade glioma (LGG) dataset. We first examined DNA methylation as a direct functional readout of D-2HG activity in IDH-mutant tumors, using 1,308 IDH-mutant-sensitive CpG probes identified from the ∼450,000 HM450 array (*4*). Whereas expression of *SWELL1*, the essential subunit shared by all VRAC complexes, did not significantly correlate with CpG methylation across the IDH-mutant cohort, expression of *LRRC8C*, the channel component that confers D-2HG transport specificity, significantly stratified mean CpG methylation across these probes and was inversely correlated with mean β-values. (**Fig. 5K** and **5L**). Consistent with this, *LRRC8C* expression was also inversely correlated with mean methylation across another set of 131 CpG probes that distinguish G-CIMP-high from G-CIMP-low tumors (*4*) (**Fig. S7E**).

Importantly, the *LRRC8C* CpG island promoter was constitutively unmethylated and did not differ between *LRRC8C*-high and *LRRC8C*-low tumors (mean methylation: 2.19% versus 2.25%, p=0.072) (**Fig. S7F**), ruling out the possibility that *LRRC8C* expression is itself suppressed by G-CIMP-driven promoter hypermethylation. Instead, these findings argue against *LRRC8C* being a passive downstream target of G-CIMP-driven silencing and are consistent with *LRRC8C* expression acting upstream of the tumor epigenetic state. Consistent with this interpretation, higher *LRRC8C* expression positively correlated with expression of canonical G-CIMP-silenced tumor suppressor genes, including *MGMT*, *CDKN2A*, *RB1*, *MLH1*, *CDKN2B*, *RASSF1*, *DAPK1*, and *CDH1* (**Fig. S7G**), indicating reduced epigenetic silencing at these loci in *LRRC8C*-high tumors. Moreover, higher *LRRC8C* expression positively correlated with EMT gene signatures and SOX2 transcriptional target activity in the IDH-mutant tumors (**Fig. 5M** and **S7H**), consistent with our ATAC-seq and RNA-seq findings in primary mouse gliomasphere cells (**Fig. 3** and **S5**). We also evaluated *SWELL1* and *LRRC8C* expression across molecularly defined IDH-mutant glioma groups. Tumors were grouped according to available TCGA molecular annotations as IDH-mutant, 1p/19q-codeleted oligodendroglioma (grade 2, n = 91; grade 3, n = 72) or IDH-mutant, 1p/19q-non-codeleted astrocytoma (grade 2, n = 131; grade 3, n = 119). *LRRC8C* expression was lower in 1p/19q-codeleted oligodendroglioma than in astrocytoma at both grade 2 and grade 3, and expression increased with grade within astrocytoma (grade 3 versus grade 2). In contrast, *SWELL1* expression was modestly higher in oligodendroglioma than in astrocytoma, but showed no grade-associated differences within either entity (**Fig. S7I**).

Given the established role of D-2HG in TME immunosuppression, we next examined the relationship between *LRRC8C* expression and immune landscape in IDH-mutant tumors. Estimating the Proportion of Immune and Cancer cells (EPIC) deconvolution analysis revealed that *LRRC8C*-high tumors harbored modestly increased fractions of macrophages, CD4^+^ T cells, and endothelial cells, accompanied by a corresponding reduction in tumor cell fraction (**Fig. S7J**). To assess the functional state of this immune infiltrate, we reassessed the IDH-mutant RNA-seq data and correlated *LRRC8C* expression with a composite immunosuppression score derived from ranked expression of *TGFB2*, *IL10*, *IDO1*, *ARG1*, *CD274*, *IKZF2*, *HAVCR2*, and *CTLA4*. *LRRC8C*expression positively correlated with this immunosuppression score across IDH-mutant tumors, and this association persisted after adjustment for tumor purity (**Fig. 5N**). To further resolve the immunosuppressive landscape, we performed purity-adjusted partial Spearman correlation analysis using 27 marker genes spanning five immunosuppressive categories. Notably, *LRRC8C* expression positively correlated with checkpoint ligands (*PD-L1*, *PD-L2*, *CD80*, *CD86*), co-inhibitory receptors (*CTLA-4*, *TIM-3*), the Treg marker Helios (*IKZF2*), and suppressive cytokines and myeloid immunosuppression markers (*TGFB2, COX-2, IL-10, VEGFA*, *IDO1, ARG1, S100A9, S100A8, IL-8*, and *iNOS*) (**Fig. 5O**), collectively consistent with a broadly immunosuppressive microenvironment in *LRRC8C*-high tumors. Together, these findings support a role for LRRC8C-containing VRAC in promoting D-2HG-mediated immunosuppression in IDH-mutant gliomas.

To examine the prognostic significance of *LRRC8C* expression, we performed genome-wide multivariate Cox regression analysis across 17,639 expressed genes in patients with IDH-mutant LGG, adjusting for tumor grade, age, and tumor purity. *LRRC8C* emerged as the top-ranked gene (104th of 17,639; top 0.6%) significantly associated with poor overall survival (hazard ratio (HR) = 6.64; p = 7.49E-05) (**Fig. 5P**). Kaplan-Meier analysis further demonstrated that patients with high *LRRC8C* expression exhibited markedly reduced survival, with a median overall survival of 78.2 months compared with 145 months in *LRRC8C*-low patients (**Fig. 5Q**). Together, these clinical data identify *LRRC8C* expression as a correlate of reduced epigenetic silencing, enhanced immunosuppression, and poor prognosis in human IDH-mutant gliomas.

### Pharmacological VRAC inhibition phenocopies SWELL1 deletion and synergizes with immune checkpoint blockade in a mouse model of IDH-mutant glioma

Building on our genetic studies, we next sought pharmacological proof of concept for targeting VRAC in IDH-mutant gliomas. Drug repurposing provides a cost-effective strategy to rapidly translate mechanistic insights into therapeutic approaches. Dicumarol, a clinical anticoagulant, was recently identified as a potent VRAC inhibitor (*23, 59*). In mouse mIDH gliomasphere cells, dicumarol treatment reduced D-2HG release, increased intracellular D-2HG accumulation, and suppressed cell proliferation, phenocopying genetic deletion of SWELL1 (**Fig. 6A** and **6B**). Similar anti-proliferative effects were also observed in dicumarol-treated human mIDH glioma cells (**Fig. S8A**). Notably, dicumarol had no effect on proliferation of mouse wtIDH or mIDH SKO cells (**Fig. S8B**), supporting its on-target activity in vitro. Because dicumarol exhibits limited blood–brain barrier permeability, we delivered the drug intracerebrally to inhibit VRAC activity, as previously validated in a mouse experimental stroke model (*59*). To enable dicumarol delivery, we implanted a dialysis cannula (*60*), a route that confines exposure largely to the cranial cavity and minimizes systemic effects, at the time of intracranial injection of mouse mIDH gliomasphere cells. Local intracerebral dicumarol delivery was previously shown not to cause excessive hemorrhage in treated brain hemispheres (*59*), and in this study cannula implantation with three weeks of dicumarol treatment produced no weight loss or neurological deficits in tumor-free mice. Dicumarol treatments were initiated one week after tumor implantation and administered three times weekly for three weeks. Notably, dicumarol prolonged survival in mice bearing mIDH glioma (median survival [MS]: vehicle, 24 days; dicumarol, 29 days) (**Fig. 6C**), comparable to the survival benefit observed with SWELL1 deletion (MS: 31.5 days) (**Fig. S8C**). FACS profiling revealed that dicumarol treatment increased intratumoral CD8⁺ T cell abundance and enhanced the effector and activation phenotype in both CD8⁺ and CD4⁺ T cells, as evidenced by elevated IFNγ, granzyme B, surface CD107a, IL-2, Ki67, and TNFα expression, along with reduced PD-1 expression (**Fig. 6D–F** and **Fig. S8D–F**). This closely mirrored the effects observed following genetic deletion of SWELL1 (**Fig. 2E–L**). Importantly, dicumarol conferred neither survival benefit (MS: 29.5 days) nor enhanced intratumoral T cell effector/activation marker expression in mice bearing mIDH SKO gliomas (**Fig. 6D–F** and **Fig. S8C–F**). This genetic epistasis suggests that the *in vivo* activity of dicumarol requires tumoral VRAC and argues against confounding by anticoagulant or other off-target effects. Together, these results demonstrate that pharmacological inhibition of VRAC suppresses tumor growth, restores anti-tumor immunity, and prolongs survival in a mouse model of mIDH glioma.

**Figure 6.**
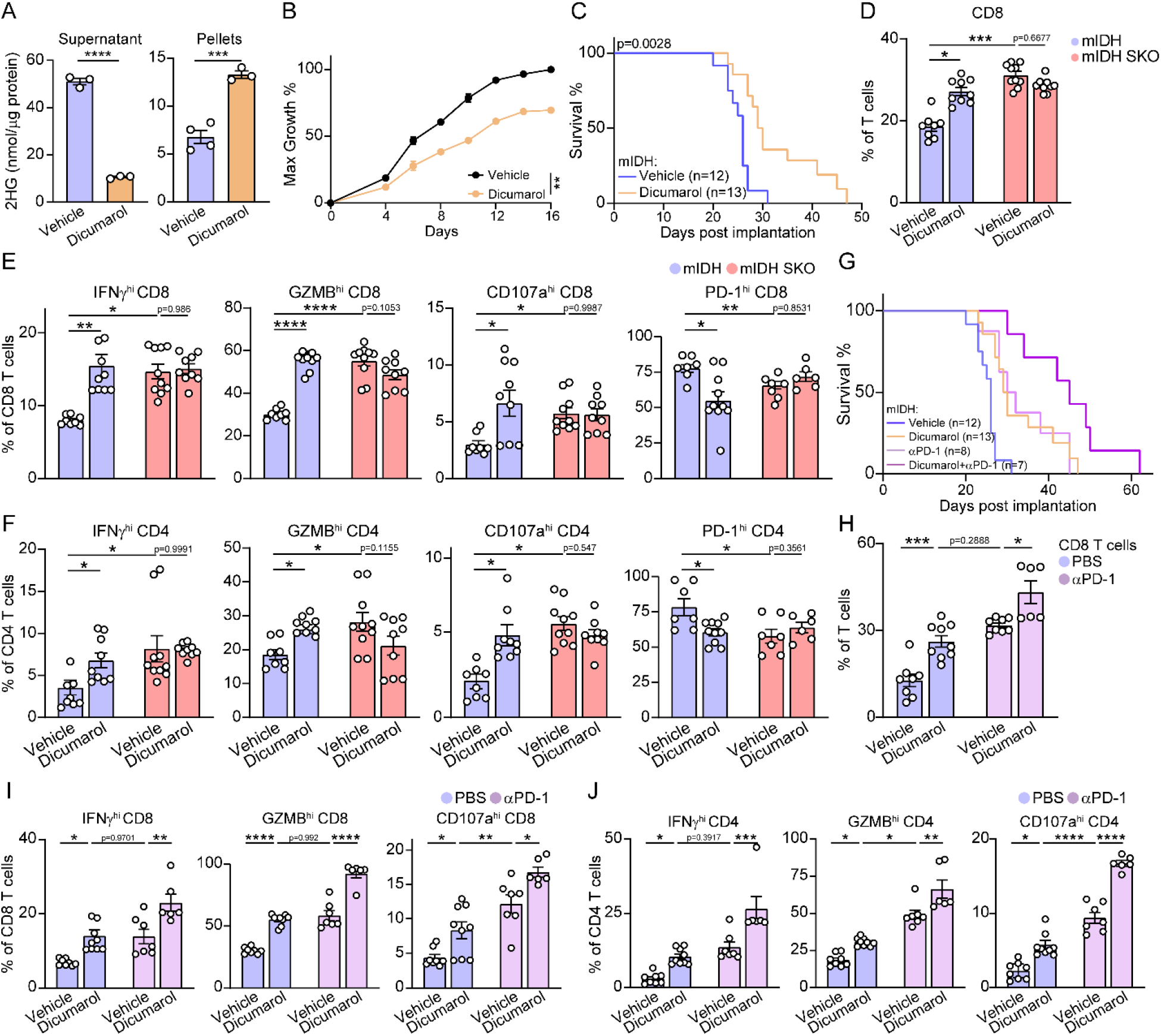
Pharmacological VRAC inhibition phenocopies SWELL1 deletion and synergizes with immune checkpoint blockade in a syngeneic mouse model of IDH-mutant glioma. **A**, 2HG quantification in cell culture supernatants (left) or pellets (right) from mouse gliomasphere cells (n = 3 to 4). Cells were treated with dicumarol (20 μM) for > 7 days before 48 h in culture for sample collection. **B**, Colony formation assay for mouse gliomasphere cells treated with 20 μM dicumarol (n = 3). Data normalized to the maximum absorbance within the comparison. **C**, Survival of mice implanted with 7 x 10^4^ mouse gliomasphere cells treated with dicumarol. Data shared with (**G**). **D,** FACS quantification of CD8+ T cell populations in mouse gliomasphere tumors treated with dicumarol (n = 8 to 10). mIDH vehicle and dicumarol are shared with (**H**). **E–F**, FACS quantification of CD8+ T (**E**) or CD4+ T cell (**F**) populations for high IFNγ, Granzyme B, surface CD107a, and PD-1 populations in mouse gliomasphere tumors treated with dicumarol (n = 6 to 7). GZMB, granzyme B. mIDH vehicle and dicumarol are shared with (**I–J**). **G**, Survival of mice implanted with 7 x 10^4^ mouse gliomasphere cells treated with dicumarol and anti-PD1 antibody. αPD-1, anti-PD1 antibody. mIDH vehicle and dicumarol are shared with (**C**). **H,** FACS quantification of CD8+ T cell populations in mouse gliomasphere tumors treated with dicumarol and anti-PD1 antibody (n = 6 to 10). αPD-1, anti-PD1 antibody. PBS vehicle and dicumarol are shared with (**D**). **I–J**, FACS quantification of CD8+ T (**I**) or CD4+ T cell (**J**) populations for high IFNγ, Granzyme B, and surface CD107a populations in mouse gliomasphere tumors treated with dicumarol and anti-PD1 antibody (n = 6 to 10). GZMB, granzyme B. αPD-1, anti-PD1 antibody. PBS vehicle and dicumarol are shared with (**E–F**). Data are reported as mean ± SEM. Unpaired t-test for **A.** Two-way ANOVA with Sidak’s test for **B, D, E, F, H, I, J.** Mantel-Cox test for **C, G.** *p < 0.05, **p < 0.01, ***p < 0.001, and ****p < 0.0001.

Given the role of D-2HG in suppressing the function of tumor-infiltrating immune cells, we next evaluated whether VRAC inhibition could enhance the efficacy of immune checkpoint blockade. While monotherapy with either dicumarol or anti-PD-1 antibody extended survival to a similar extent (MS: vehicle, 24 days; dicumarol, 29 days; anti-PD-1, 30 days), combination therapy produced a striking synergistic benefit, extending MS to 45 days (**Fig. 6G**). Mice receiving combined treatment exhibited a robust expansion of tumor-infiltrating CD8⁺ T cells, accompanied by an increased effector/activation phenotype in both CD8⁺ and CD4⁺ T cell populations (**Fig. 6H– J** and **Fig. S8G**). Together, these findings demonstrate that pharmacological inhibition of VRAC not only relieves immunosuppression but also sensitizes mIDH gliomas to immune checkpoint blockade, providing preclinical proof of concept for a combinatorial therapeutic strategy.

## Discussion

Here we identify SWELL1/LRRC8C-containing VRAC as the principal export pathway for the key oncometabolite D-2HG. Although several transporters have been proposed to mediate D-2HG transport (*16–19, 61*), our functional screen and validation studies identify VRAC as a dominant and broadly utilized conduit for D-2HG export from IDH-mutant cells. The large pore of this anion channel makes it particularly well suited to mediate the release of negatively charged D-2HG, which accumulates to millimolar intracellular concentrations (*7–9*). While previous studies have shown that tumor-derived D-2HG suppresses T cell function *in vitro* and correlates with immunosuppression in patients with IDH-mutant glioma (*11, 12*), direct *in vivo* evidence for this mechanism has been lacking. Identification of VRAC as the principal D-2HG release pathway, together with the use of immunocompetent mouse glioma models, enabled testing of this hypothesis *in vivo*. Both genetic deletion of *SWELL1* and pharmacological inhibition of VRAC reprogrammed the TME toward an immunoactive state, establishing VRAC-mediated D-2HG export as a key mechanistic link between tumor metabolism and immune evasion in IDH-mutant gliomas. Notably, extracellular D-2HG has also been implicated in the high incidence of preoperative epilepsy in patients with IDH-mutant glioma by acting as a glutamate mimic at NMDA receptors and enhancing neuronal excitability (*62*). Future studies will investigate whether VRAC-dependent D-2HG release contributes to glioma-associated epileptogenesis.

Human mIDH gliomas typically progress more slowly and are associated with significantly better clinical outcomes than wtIDH tumors of the same histological grade (*32, 36–39*). Clinical and mechanistical studies suggest that D-2HG plays a paradoxical role in glioma biology: it drives cellular transformation during tumor initiation (*5, 6*) while simultaneously imposing metabolic and epigenetic constraints that can limit proliferation (*32, 36–38, 53–56*). This duality may help explain why the recently approved mIDH inhibitor vorasidenib shows its greatest therapeutic benefit in early-stage, non-contrast-enhancing gliomas (*63*). It also suggests that IDH-mutant cancer cells may actively clear this oncometabolite to sustain cell growth, although such mechanisms have remained unknown. While D-2HG dehydrogenase can metabolize D-2HG generated as a metabolic by-product in normal cells (*64*), its mitochondrial localization likely limits its ability to eliminate the large cytosolic pools of D-2HG produced by mutant IDH1 (*41*). VRAC-mediated D-2HG efflux therefore represents a previously unrecognized D-2HG clearance pathway in mIDH cells. This export mechanism prevents D-2HG overload, thereby mitigating metabolic stress, reducing reliance on compensatory metabolic pathways, and restraining excessive histone and DNA hypermethylation to preserve pro-proliferative transcriptional programs. Consistent with this epigenetic framework, patients with IDH-mutant gliomas exhibiting high levels of the glioma CpG island methylator phenotype (G-CIMP-high) have significantly better overall survival than G-CIMP-low patients (*4, 40*). By exporting D-2HG, VRAC serves two critical functions: alleviating intracellular D-2HG-induced cellular stress while simultaneously suppressing immunosurveillance in the tumor microenvironment. Consistent with this model, pharmacological inhibition of VRAC suppresses mIDH cell proliferation, enhances anti-tumor immunity, synergizes with immune checkpoint blockade, and prolongs survival in glioma-bearing mice. The definitive translational advancement of this pathway will require brain-penetrant, selective VRAC inhibitors, which are not yet available. Our study therefore provides proof of principle for therapeutic targeting of this pathway rather than an immediately translatable regimen.

Our data indicate that VRAC blockade restrains mIDH glioma growth through two mechanistically distinct arms. The first is tumor-extrinsic: reduced D-2HG release alleviates immunosuppression in the TME, as reflected by myeloid activation and enhanced CD8^+^ and CD4^+^ T cell effector function (**Fig. 2E–L**). The second is tumor-intrinsic: intracellular D-2HG accumulation drives epigenetic remodeling and mitochondrial metabolic stress, directly limiting tumor cell proliferation (**Fig. 3–5**). These tumor-intrinsic effects were sufficient to extend survival in immunodeficient *SCID* hosts, which lack functional T and B cells (**Fig. 5I–J**). Together, both mechanisms make important contributions to host survival. Future dedicated cell-type-specific perturbation studies will be needed to define the sensitivity of individual TME populations to tumor-derived D-2HG and to dissect how tumor-extrinsic immune remodeling and tumor-intrinsic metabolic and epigenetic stress each contribute to VRAC blockade–mediated tumor control.

Among LRRC8 paralogues, LRRC8C appears to play an important role in mediating D-2HG transport, consistent with prior studies implicating this subunit in glutamate permeation (*65*), a structurally related metabolite. However, the effect of *LRRC8C* knockdown on D-2HG release was more modest than that observed following *SWELL1* knockdown, suggesting that additional LRRC8 subunits may also contribute to D-2HG export. In particular, LRRC8E shares overlapping substrate selectivity with LRRC8C and may partially compensate for its loss (*66*). For this reason, we focused our genetic studies on *SWELL1*, whose deletion abolishes all VRAC channel activity and produces a stronger phenotype. Interestingly, despite the more robust phenotype associated with *SWELL1* deletion experimentally, expression of *LRRC8C*, but not *SWELL1*, strongly correlated with reduced DNA methylation, elevated EMT and SOX2 transcriptional programs, immunosuppressive gene signatures, and poor survival in the IDH-mutant TCGA-LGG cohort. This distinction likely reflects the different biological roles of these channel components: SWELL1 functions as the obligate subunit shared by all VRAC complexes, whereas LRRC8C may more specifically mark the subset of VRAC channels specialized for D-2HG transport. Accordingly, in bulk RNA-seq tumor datasets, *LRRC8C* expression may more accurately reflect the functional output of D-2HG-permeable VRAC activity in human tumors. More broadly, these findings highlight the complexity of VRAC regulation in cancer biology. In addition to subunit composition, VRAC activity is dynamically regulated by osmotic stress and diverse extracellular and intracellular stimuli, including reactive oxygen species, ATP, and sphingosine-1-phosphate (*23, 26, 67*). Moreover, we recently identified puromycin-sensitive aminopeptidase (PSA) as an auxiliary subunit that binds the cytosolic LRR domains of SWELL1 and suppresses basal VRAC activity (*68*). PSA also regulates VRAC-mediated transport of the anti-cancer drug cisplatin (*69*) and the innate immune messenger cGAMP (*68*), raising the possibility that PSA similarly modulates D-2HG export in mIDH glioma cells.

Our findings may have broader implications beyond gliomas, as IDH mutations and D-2HG accumulation occur in multiple cancer types, including cholangiocarcinoma, chondrosarcoma, and acute myeloid leukemia (AML). In these cancers, intracellular D-2HG drives epigenetic hypermethylation and impairs cellular differentiation, similar to its effects in IDH-mutant gliomas (*70–74*). Metabolically, IDH mutations disrupt the TCA cycle, enhance glutaminolysis, and induce mitochondrial stress (*52–56, 75, 76*). This metabolic rewiring has also been implicated in conferring resistance to mIDH inhibitors in AML (*75*). Beyond these cell-intrinsic effects, IDH mutations suppress anti-tumor immunity in cholangiocarcinoma (*77*) and chondrosarcoma (*78*), mirroring the immunosuppressive TME observed in IDH-mutant gliomas. Notably, the high levels of D-2HG observed in these malignancies suggest active release from tumor cells, and VRAC is ubiquitously expressed across tissues (*20, 22*). Together, these observations raise the testable possibility that VRAC-dependent D-2HG transport may also shape the metabolic state and immune microenvironment of non-CNS IDH-mutant malignancies. However, direct functional evidence in these disease contexts is currently lacking. Thus, VRAC-mediated D-2HG export may represent a conserved adaptive mechanism across IDH-mutant cancers, enabling tumor cells to simultaneously maintain cellular fitness while evading immune surveillance.

## Acknowledgements

We thank Dr. Shuli Xia at Kennedy Krieger Institute and Dr. Timothy Chan at Memorial Sloan Kettering Cancer Center for providing mIDH knock-in U87 cells and patient-derived TS603 cells, respectively. We thank Drs. Steven Claypool and Nanami Senoo for the technical assistance on Seahorse analysis, Dr. Xinzhong Dong and Aleksander Geske for the technical assistance on Flow cytometry, Dr. Santosh Yadav for the assistance on mouse MRI imaging, and Dr. Ruchita Kothari and the members of Qiu lab for valuable discussion.

This work was supported by National Institutes of Health grant R35GM124824, R01NS118014, RF1NS134549, McKnight Scholar Award, Klingenstein-Simon Scholar Award, Sloan Research Fellowship in Neuroscience, Randall J. Reed Scholars Award and American Heart Association Established Investigator Award to Z.Q.

## Data Availability Statement

ATAC-seq and RNA-seq raw data are deposited on GEO database (GSE345819, GSE345834, and GSE345835). Supporting Data Values for all graphs are provided as the accompanying Supplementary Raw data. mIDH-transduced WT and SWELL1 KO MC38 and HeLa cells as well as SWELL1 KO mouse gliomasphere, SJGBM2, TS603, and U87 cells were generated in this study. All will be made available from the lead contact upon request. Further information and requests for resources should be directed to and will be fulfilled by the Lead Contact, Zhaozhu Qiu.

## Author contributions

H.Y.C. and Z.Q. designed the study; H.Y.C, L.M., J.H., L.C., J.C., W.L., R.Z., and L.K. performed research; H.Y.C., L.M., L.C., J.C., Y.L., Y.Y., L.K., S.S., J.Z., L.H., and Z.Q. contributed to data analysis and interpretation; W.Z., and M.G.C. contributed research material; H.Y.C. and Z.Q. wrote the manuscript with inputs from all authors.

## Materials and Methods

### Mice

The use of all mice in the study was approved by the Johns Hopkins University Animal Care and Use Committee. All mice were housed in a specific pathogen free (SPF) environment, under standard 12-hour light/12-hour dark cycle with *ad libitum* access to food and water. C57BL/6J (000664) and *SCID* (B6.Cg-*Prkdc*^scid^/SzJ; 001913) were purchased from The Jackson Laboratory. Adult male mice between 8-12 weeks of age were used for experiments as specified with age-and sex-matched littermates as controls.

### Cell Culture

The primary mouse gliomasphere cells were generated from a *de novo* mouse glioma model (*32*) carrying shRNA against ATRX and TP53 (referred to wtIDH cells) while mIDH cells were generated by introducing mutant IDH1^R132H^ to wtIDH cells (*32*). Cells were maintained in neural stem cell medium [DMEM/F12 (Gibco) with L-Glutamine (Gibco, 11320033), B-27 supplement (Gibco, 12587010), N-2 supplement (Gibco, 17502048), penicillin-streptomycin (Gibco, 15140122), and Normocin (InvivoGen, ant-nr-1)] at 37°C in humidified 5% CO2 incubator. Recombinant FGF (Peprotech, 100-18B) and EGF (Peprotech, AF-100-15) were supplemented at 20 ng/mL in the neural stem cell medium upon cell culture. Mouse gliomasphere cells were maintained below 10 passages for all experiments.

Patient-derived TS603 cell is a WHO grade III anaplastic oligodendroglioma with 1p/19q co-deletion and IDH1^R132H^ mutation. TS603 cells were generously provided by Dr. Timothy Chan (Memorial Sloan Kattering Cancer Center). Cells were maintained as floating spheres in Neurobasal medium (Gibco, 21103049) supplemented with GlutaMAX (Gibco, 35050061), B-27 supplement (Gibco, 12587010), N-2 supplement (Gibco, 17502048), penicillin-streptomycin (Gibco, 15140122), and Normocin (InvivoGen, ant-nr-1)] at 37°C in humidified 5% CO2 incubator. Recombinant FGF (Peprotech, 100-18B) and EGF (Peprotech, AF-100-15) were supplemented at 20 ng/mL in the medium upon cell culture. TS603 cells were maintained below 10 passages for all experiments.

Monoallelic mIDH U87 human glioblastoma cell line was generated by CRISPR knock-in (*15*) and generously provided by Dr. Shuli Xia (Kennedy Krieger Institute). Cells were maintained in Dulbecco’s modified Eagle’s medium (DMEM) supplemented with L-Glutamine (Gibco, 11965092), 10% heat inactivated (HI)-FBS, penicillin-streptomycin (Gibco, 15140122) at 37°C in humidified 5% CO2 incubator. mIDH-transduced SJGBM2 pediatric glioma cells were generated as previously described (*32*) and maintained in improved minimum essential medium (IMEM) supplemented with L-Glutamine (Gibco, A1048901), 20% HI-FBS, penicillin-streptomycin (Gibco, 15140122) at 37°C in humidified 5% CO2 incubator.

MC38 mouse colon adenocarcinoma cells were purchased from Millipore Sigma (SCC172). Cells were maintained in RPMI 1640 medium supplemented with L-Glutamine (Gibco, 11875085), 10% HI-FBS, penicillin-streptomycin (Gibco, 15140122) at 37°C in humidified 5% CO2 incubator. HeLa cells were maintained in DMEM supplemented with L-Glutamine (Gibco, 11965092), 10% HI-FBS, penicillin-streptomycin (Gibco, 15140122) at 37°C in humidified 5% CO2 incubator.

### Gene Transduction and CRISPR Gene Deletion

SWELL1 knockout (SKO) cells were generated by CRISPR-Cas9 technology. For mouse gliomasphere cells, guide RNA1 (GCTGTGTGTCCGCAAAGTAG) or guide RNA2 (TGATGATTGCTGTCTTTGGA) targeting mouse *Swell1/Lrrc8a* was cloned into LentiCRISPR-v2-PuroR plasmid (Addgene #98290, a gift from Dr. Feng Zhang). Lentiviral particles containing mouse *Swell1* sgRNA were packaged using the third-generation lentiviral system. 24 hours after viral transduction, the medium was changed to fresh culture medium. Puromycin 10 µg/mL was used to select successfully transduced cells. For U87, SJGBM2, and TS603 cells, guide RNA (TGATGATTGCCGTCTTCGGG) targeting human *SWELL1/LRRC8A*, was cloned into PX458-mCherry, a modification of the plasmid pSpCas9(BB)-2A-GFP (PX458) (Addgene #48138, a gift from Dr. Feng Zhang). For MC38 and HeLa cells, mouse *Swell1* guide RNA1 or human *SWELL1* guide RNA, was cloned into PX458-GFP plasmid (Addgene #48138, a gift from Dr. Feng Zhang), respectively. 48-72 hours after transfection, the medium was changed to fresh culture medium. For U87, SJGBM2, TS603, MC38, and HeLa cells, cells post-transfection were then FACS sorted for positive GFP or mCherry signal. SWELL1 protein expression for all KO cells was validated using immunoblotting.

For the generation of mIDH MC38 and HeLa cells, cells were co-transfected with pCMV(CAT)T7-SB100 (Addgene #34879) and pKT-IDH1(R132H)-IRES-Katushka (Addgene #124257) plasmids at 1:10 ratio. 48 hours after transfection, the medium was changed to fresh culture medium. Cells transfected were then FACS sorted for positive Katushka signal to select successfully transduced cells. IDH1^R132H^ protein expression was validated using immunoblotting.

### 2HG#Measurements

2-HG levels were quantified using GC–MS. To 1 mL of medium or cell lysate, 0.4 μg of ^13^C_5_-DL-2-hydroxyglutaric acid, disodium salt (Cambridge Isotopes) was added, samples were acidified with 3N HCl, then extracted twice with 2 mL of ethylacetate. Pooled solvent layers were evaporated to dryness under nitrogen and the residues derivatized with BSTFA+ 1% TMCS. For SIM-GC/MS analysis, 1 µL was injected into an Agilent 8890/5977B GC/MSD system utilizing an Agilent DB5-ms 25 meter, 0.25 mm ID capillary column. The GC oven program was set as follows: 70°C for 2 minutes.; 4°C/min ramp to 180°C; 30°C/min ramp to 290°C; hold for 10 minutes. Mass spectrometric data were collected in the selected ion mode at *m/z = 247.2* for native 2-hydroxyglutarate and *m/z = 251.1* for ^13^C_5_-2-hydroxyglutaric acid.

For the VRAC-mediated D-2HG release under hypotonic stimulation, U87 mIDH and mIDH SKO cells were seeded in 12-well plates at 10⁵ cells/well. On the day of the experiment, the culture medium was replaced with a hypotonic bath solution (90 mM NaCl, 2 mM KCl, 1 mM MgCl₂, 2 mM CaCl₂, 10 mM HEPES, 10 mM glucose; pH adjusted to 7.3 with NaOH; osmolality ∼212 mOsm/kg, without mannitol) and cells were incubated at 37°C for 30 minutes. Following incubation, supernatants were collected and centrifuged at 12,000 ×g for 10 minutes at 4°C to remove cellular debris before 2HG quantification by GC-MS as described above.

For D-2HG measurements in mouse tissues, brain samples (20-100 mg) were homogenized in 0.1N salt-saturated HCl, then extracted as above. O-acetylated di-(-)-2-butyl ester derivatives were prepared as previously described(*79*) to allow for separation of D and L isomers of 2-hydroxyglutarate by GC–MS. Mass spectrometric data were collected in the selected ion mode at *m/z = 173* for native 2-hydroxyglutarate and *m/z = 178* for ^13^C_5_-2-hydroxyglutarate.

### ATP Depletion

U87 cells were seeded to 12-well plates at 10^5^ cells/well density and maintained in serum-free ATP depletion medium (no glucose DMEM (Gibco, A1443001) with penicillin-streptomycin (100 U/mL), 6 mM 2-deoxy-D-glucose, and 5 mM NaN3 for 4 hours before the supernatants were collected for 2HG quantification.

### Molecular Weight Cutoff (MWCO) Filtration

To determine whether extracellular 2HG is released as a freely soluble molecule rather than in association with extracellular vesicles or proteins, cell culture supernatants from U87 mIDH cells were subjected to 10-kDa molecular weight cutoff (MWCO) filtration using Amicon Ultra-0.5 centrifugal filter units (Millipore, #UFC501024). Samples were centrifuged at 8,000 ×g for 10 minutes at 4°C. The flow-through fraction was collected and used for 2HG quantification by GC-MS as described above.

### LDH Assay

U87 cells were seeded to 12-well plates at 10^5^ cells/well density, and the supernatants were collected at indicated time points for both 2HG quantification and LDH activity assay. 100 uL of supernatant was used for the LDH assay using CyQUANT LDH Cytotoxicity Assays (Invitrogen) per manufacturer’s instructions.

### siRNA Transfection

U87 cells were seeded to 12-well plates at 5 × 10^4^ cells/well density or 6-well plates at 5 × 10^5^ cells/well density 12-18 hours before the siRNA transfection. Cells were then transfected with Dharmacon ON-TARGETplus SMARTPool siRNA containing 4 siRNAs targeting each gene (details in Supplemental Material Table) at 50 nM using Lipofectamine RNAiMAX transfection reagent (Invitrogen) for 48 hours. As controls, scrambled siRNA duplexes (QIAGEN) were used. After the transfection, cell culture supernatant was replaced by fresh cell culture medium before collection for 2HG quantification.

### Patch Clamp Electrophysiology

Whole-cell patch clamp recordings were performed as previously described (*23*). For hypotonicity-activated VRAC current recordings, whole-cell patch-clamp configuration was established in an isotonic bath solution containing 90 mM NaCl, 2 mM KCl, 1 mM MgCl_2_, 2 mM CaCl_2_, 10 mM HEPES, 10 mM glucose, and 100 mM mannitol (pH adjusted to pH 7.3 with NaOH and osmolality adjusted to 312 mOsm/kg), and then a hypotonic solution that has the same ionic composition but without mannitol was applied. Recording electrodes (2 to 4 megohms) were filled with a standard internal solution containing 133 mM CsCl, 10 mM HEPES, 2 mM CaCl2, and 5 mM EGTA (pH was adjusted to 7.2 with CsOH, and osmolality was 290 to 300 mOsm/kg).

To determine the D-2HG permeability of VRAC, recording electrodes (2 to 4 megohms) were filled with the Cl^−^-based intracellular solution containing 90 mM NaCl, 10 mM HEPES, and 100 mM mannitol (pH adjusted to pH 7.3 with NaOH and osmolality adjusted to 310 mOsm/kg). D-2HG^2−^-based intracellular solution was made by replacing 90 mM NaCl and 100 mM mannitol with 90 mM Na_2_D-2HG^2−^ to maintain the same osmolality. The reversal potentials were determined using the ramp protocol. Relative permeability of D-2HG^2−^ to Cl^−^ was estimated on the basis of the Goldman-Hodgkin-Katz flux equation.

### Orthotopic Mouse Glioma Models

Male C57BL/6J mice, aged 8-10 weeks old, were used for the syngeneic orthotopic glioma model. The primary mouse gliomasphere cells used for orthotopic implantation were derived from a de novo glioma model induced in C57BL/6 mice (*32*) and are therefore syngeneic to C57BL/6J hosts; these tumors are referred to as syngeneic orthotopic gliomas throughout. Intracranial tumors were established by stereotactic injection of 5 × 10^4^ primary mouse gliomasphere cells into the right striatum using a 22-gauge Hamilton syringe (1 μL per minute) with the following coordinates: +1.00 mm anterior, 2.5 mm lateral, and 3 mm deep.

For TS603 human glioma orthotopic xenograft model, male *SCID* mice aged 8-10 weeks were used. Intracranial tumors were established by stereotactic injection of 1 × 10^5^ cells with the same specifications as mouse gliomasphere orthotopic xenograft model.

For dicumarol treatments, intracranial tumors were established by stereotactic injection of 7 × 10^4^ primary mouse gliomasphere cells, following the same specifications as described above. Immediately after tumor cell injection, a microdialysis cannula (*60*) (RWD) was implanted at the injection site and secured to the skull using miniature screws (RWD) and dental cement (Shofu). Following surgery, mice were recovered in a warmed cage and monitored for stress or neurological symptoms until fully recovered. Cannula implantation alone did not induce weight loss or neurological deficits in tumor-free mice over a three-week post-surgical period. Dicumarol (in 2 μL PBS, 500 μM, 0.33 % DMSO) was administered via the cannula at a rate of 1 μL/min starting one week post-surgery, three times per week for three weeks. For αPD-1 immune blockade, 100 μg of antibody (Bio X cell, BE0033-2) in 200 μL PBS was injected intraperitoneally on day 9 and 16 post-surgery.

### Magnetic Resonance Imaging (MRI)

T1-weighted MRI imaging with gadolinium-based contrast agent in mice bearing orthotopic gliomas were performed at MRB Molecular Imaging Service Center at Johns Hopkins University School of Medicine. Specifically, mice underwent an imaging session under 1-3% isoflurane inhalation. Animals were placed in a heated, MRI-compatible cradle with continuous monitoring of respiration and body temperature. An FDA-approved gadolinium-based contrast agent was administered intravenously via the tail vein at 0.05–0.1 mmol/kg diluted with sodium chloride (1:3) as a single bolus 5 minutes before imaging. Each imaging session included both pre-and post-contrast T1 scans using a 9.4T small-animal MRI system with optimized TR/TE parameters (e.g., 3,500 ms TR, 30 ms TE). Total scan time per session was 5 minutes, with total anesthesia duration not exceeding 8 minutes. Following imaging, mice were recovered in a warmed cage and monitored until fully recover (5–15 minutes), then returned to their home cages.

### Tumor Dissociation

After euthanasia, gliomas from tumor-bearing mice were harvested after cardiac perfusion. Tumors were minced into small pieces before incubating with tumor dissociation medium containing RPMI, 1x Collagenase/Hyaluronidase (STEMCELL), and 0.15 mg/mL DNase 1 (STEMCELL) at 220-250 rpm, 37°C for 20 minutes on a shaking platform. After dissociation, 70 μm mesh strainer was used to remove undissociated tissues and ACK buffer (Quality Biological) was used to remove red blood cells before made into single cell suspension for subsequent experiments.

### Flow Cytometry (FACS) Analysis

Mice received gadolinium for MRI tumor imaging were excluded from FACS analysis. Immune cells from tumor dissociation were enriched by anti-mouse CD45 beads using magnetic cell separation (Miltenyi). FACS antibody staining was performed as previously described (*80*). In Brief, single cell suspension from tumor dissociation or cell cultures was resuspended in FACS buffer (2% FBS, 2 mM EDTA in sterile PBS). Live versus Dead cells were stained in PBS using Live/Dead Fixable Aqua Dead Cell Stain (Invitrogen). Fc receptors were then blocked by incubation with TrueStain FcX anti-CD16/32 antibody (BioLegend) at 4°C for 20 minutes, and the cells were subsequently stained with fluorophore-conjugated antibodies listed in the Supplemental Material Table, for 20 minutes at 4°C. For unconjugated primary antibodies, subsequent fluorophore-conjugated secondary antibodies were used with the same staining method. For cytokine staining, tumor-infiltrating immune cells were restimulated with 50 ng/mL PMA and 1 μg/mL ionomycin in complete ell culture medium for 4 h. Brefeldin A (BioLegend) was introduced after 1 h of incubation. Subsequently, cells were collected and subjected to surface and intracellular staining. Intracellular proteins were stained using Cyto-Fast Fix/Perm Buffer set (BioLegend), True-Nuclear Transcription Factor Buffer Set (BioLegend), or Foxp3/Transcription factor staining buffer kit (eBioscience), following manufacturer’s instructions. Live/Dead Fixable Aqua Dead Cell Stain was also used for the TS603 cell viability assay. Data was acquired on a CytoFLEX LX (Beckman Coulter) and analyzed using FlowJo (BD) or OMIQ.ai.

### DNA Methylation Assay

Fresh cell pellets were prepared and processed according to manufacturer’s instructions (Abcam). In brief, cell pellets were lysed for the genomic DNA collection before binding to the assay wells. Methylated DNA was detected by the detection antibody at room temperature for 60 minutes before incubating with Enhancer and Developer Solutions for the signal detection. After adding the Stop Solution, the antibody-bound methylated DNA was detected at 450 nm. The relative 5-mC percentage was calculated as 5-mC % = [(Sample OD -Negative Control OD) + S] / [(Positive Control OD - Negative Control OD) × 2* + P] × 100%. S refers to input positive control in ng. P refers to sample DNA in ng. *2 is a factor to normalize 5-mC in the positive control to 100%, as the positive control contains only 50% of 5-mC.

### Colony Formation and LC50 assay

For basal growth activity, primary mouse gliomasphere cells were seeded at 8,000 cells/well in 96-well plates. For αKG supplementation experiments, primary mouse gliomasphere cells were pre-treated with 10 mM αKG for at least 48 hours prior to the assay. On the day of the experiment, cells were seeded at 8,000 cells/well in 96-well plates in neural stem cell medium supplemented with 5 mM αKG to maintain supplementation throughout the assay. On the day of staining, cells were centrifuged at 300 xg for 5 minutes before fixed with 4% paraformaldehyde (PFA) for 15 minutes at room temperature and stained with 0.5% crystal violet (Millipore Sigma) for another 15 minutes. After staining, crystal violet solution was removed after centrifuged at 500 xg for 5 minutes followed by PBS wash twice. Crystal violet dyes bound to cell pellets were then solubilized by 1% ethanol and the absorbance at 570 nm was measured to represent cell colony size.

For LC50 analysis, primary mouse gliomasphere cells were seeded at 20,000 cells/well in 96-well plates 3 days before experiment. Cells were simultaneously treated with indicated chemical compounds at indicated concentrations. After 3 days, cells were centrifuged at 300 xg for 5 minutes before fixed with 4% PFA for 15 minutes at room temperature and stained with 0.5% crystal violet for another 15 minutes. After staining, crystal violet solution was removed after centrifuged at 500 xg for 5 minutes followed by PBS wash twice. Crystal violet dyes bound to cell pellets were then solubilized by 1% ethanol and the absorbance at 570 nm was measured to represent viable cells.

### ATAC Sequencing

ATAC sequencing (ATAC-seq) service was carried out by Novogene Company. In brief, cell nuclei pellets were resuspended in the Tn5 transposase reaction mix at 37°C for 30 minutes. Equimolar Adapter1 and Adapter 2 are added after transposition, and PCR is then performed to amplify the library. After the PCR reaction, libraries are purified with the AMPure beads (Beckman Coulter). Samples were then sequenced on an illumina NovaSeq 6000 System. Reads were mapped to mm10 reference mouse genome using Burrow-Wheeler Aligner (BWA) and the peak calling was performed with MACS2 algorithm. Multiple Expectation maximization for Motif Elicitation (MEME) was used for the motif prediction and the PeakAnnotator was used for the ATAC-seq peak annotations.

### RNA Sequencing

Total RNA from primary mouse gliomasphere cells was extracted using the TRIzol method (Invitrogen). RNA quality was determined by the A260/A280 and A260/A230 as well as Agilent 2100 Bioanalyzer. RNA sequencing (RNA-seq) service was carried out by Novogene Company. In brief, libraries were constructed using the NEBNext Ultra II RNA library Prep Kit for Illumina (New England Biolabs) and sequenced on an illumina NovaSeq 6000 System. Reads were mapped to GRCm39 reference mouse genome with Hisat2 v2.0.5. FeatureCounts v1.5.0-p3 was used to count the reads numbers mapped to each gene. Fragments per kilobase per million mapped reads (FPKM) of each gene was calculated based on the length of the gene and reads count mapped to this gene. For tumor intrinsic immunogenicity gene clusters, VST (variance-stabilizing transformation) normalization was applied to raw DESeq2-normalized counts using the to account for the experimental design, producing stabilized log2-scale expression values suitable for visualization and unsupervised analyses. Further differential expression analysis, Gene Set Enrichment Analysis (GSEA), Gene Ontology (GO) analysis, and KEGG pathway analysis were conducted using the DESeq2 and clusterProfiler R package. Data were visualized using GraphPad Prism and NovoMagic (Novogene).

### Integrated Analysis for ATAC-seq and RNA-seq

Integrated analysis of ATAC-seq and RNA-seq datasets was performed to identify genes exhibiting concordant changes in chromatin accessibility and transcriptional output between mIDH and mIDH SWELL1 knockout (SKO) mouse gliomasphere cells. ATAC-seq peak detection was performed per sample using a binary classification approach: a peak was considered present in a given sample if it met the significance threshold in that replicate. For the Closed+Down gene set, representing loci with reduced accessibility and decreased expression in mIDH SKO cells, candidate peaks were required to be detected in at least 2 of the control (mIDH) replicates (n_ctrl ≥ 2) and absent in all SKO replicates (n_sko = 0). Corresponding genes were further filtered by DESeq2 differential expression analysis, retaining only those with log2 fold change < -1 and adjusted p-value < 0.05 (Wald test with Benjamini-Hochberg correction), yielding 220 concordantly closed and downregulated genes. For the Open+Up gene set, representing loci with gained accessibility and increased expression in mIDH SKO cells, peaks were required to be detected in at least 2 SKO replicates (n_sko ≥ 2) and absent in all control replicates (n_ctrl = 0), with corresponding DESeq2 log2 fold change > +1 and adjusted p-value < 0.05, yielding 175 concordantly opened and upregulated genes. The resulting set of overlapping genes was subjected to Gene Ontology enrichment analysis. Finally, representative loci were visualized as signal tracks using the Integrative Genomics Viewer (IGV) aligned to the mm10 reference genome.

### TCA cycle and Glutaminolysis Metabolomics

Targeted metabolomic quantification for TCA cycle, glutaminolysis, glycolysis, and purine metabolites was performed using complementary Agilent 8890 GC system coupled with a 7010 Triple Quadrupole mass spectrometer (GC–MS) or Thermo Vanquish ultra-high-performance liquid chromatography (UHPLC) system coupled to a Q Exactive Hybrid Quadrupole-Orbitrap mass spectrometer (LC–MS) platforms. GC–MS was employed to quantify a broad panel of organic acids, amino acids, and purine metabolites following derivatization-based analysis, while LC–MS was used for sensitive, targeted quantification of key polar and redox-related metabolites, including TCA intermediates. Metabolite abundances were normalized to protein content. Raw GC–MS and LC–MS data were processed using Agilent MassHunter Quantitative analysis and Xcalibur Quan Browser with batch reprocessing, respectively. A predefined processing method was applied to extract peak areas of targeted metabolites. All chromatographic peaks were visually inspected to confirm correct peak assignment, particularly in cases of retention time shifts due to column variability. Manual peak integration was performed when necessary to ensure accurate quantification, and verified peak areas were exported for downstream analysis.

Adenine, citric acid, galactose, D-glucose, D-glucose-6-phosphate, fructose, fumaric acid, hypoxanthine, inosine, lactic acid, glutamine, pyruvic acid, succinic acid, uric acid, xanthine, and xanthosine were analyzed by GC–MS. Quantitative calibration curves were generated using external standards prepared at concentrations ranging from 1.25 to 50 µM. GC–MS analysis was performed using the Agilent Fiehn Library Kit per manufacturer’s instruction. Extraction solvent consisting of acetonitrile/isopropanol/water (3:3:2, v/v/v) was prepared using LC–MS–grade solvents, adjusted to neutral pH, supplemented with isotope-labeled internal standards, degassed by nitrogen bubbling for 5 minutes, and precooled to −20°C. Cell extracts were dried in a vacuum centrifuge before being reconstituted in 450 µL 50% nitrogen-degassed acetonitrile and subjected to an additional vacuum centrifuge step prior to derivatization. For chemical derivatization, dried samples were methoximated with 10 µL of methoxyamine hydrochloride (20 mg/mL in pyridine) and incubated at 30°C for 1.5 hours with shaking. Subsequently, samples were silylated by adding 91 µL MSTFA containing fatty acid methyl ester (FAME) internal markers and incubated at 37°C for 30 minutes. Derivatized extracts were transferred to glass autosampler vials with micro-inserts, capped immediately, and analyzed by GC–MS under dynamic multiple reaction monitoring (dMRM) mode. For separation, a Agilent J&W DB-5MS GC column with a 95% dimethyl/5% diphenyl polysiloxane stationary phase, measuring 30 meters in length, 0.25 millimeters in internal diameter, and coated with a 0.25-micrometer film, was used. Additionally, a 10-meter empty DuraGuard guard column was employed. 1 μL of samples was injected. The inlet temperature was 250°C. The initial oven temperature was 60°C for 1 minute, followed by a ramped of 10°C/min to 325°C and held for 12.5 minutes. The total run time was 40 minutes. Helium was used as the carrier gas at a flow rate of 1.035 mL/min. Nitrogen was used as the collision gas at 1.5 mL/min, while helium was used as the quenching gas at 4 mL/min. The MSD transfer line temperature was 270°C, the ion source temperature was 260°C, the quadrupole temperature was 150°C and a gain factor of 1 was used.

αKG, fructose-6-phosphate, and glutamate were analyzed by LC–MS. Cell pellets were homogenized in 200 µL of ice-cold 1 N perchloric acid, sonicated for 80 seconds, and centrifuged at 10,000 xg for 2 minutes at 4°C before the pH was adjusted to 8.0 using 1 M potassium hydroxide. Samples were then centrifuged again at 10,000 xg for 2 minutes at 4°C, and the resulting supernatants were used for LC–MS analysis. Calibration standards for each analyte were prepared over a concentration range of 19.5 nM to 10 µM for quantitative analysis. An Waters XBridge BEH Amide column (2.5 µm, 2.1 x 150 mm; Waters Corp., MA) was used for compound separation, with separate injections for positive and negative ionization modes. Mobile phase A was 10% acetonitrile, containing 5 mM ammonium acetate and 0.1% acetic acid, and mobile phase B was 90% acetonitrile, containing 5 mM ammonium acetate and 0.1% acetic acid. For αKG, fructose-6-phosphate, and glutamate analysis in negative mode, isocratic elution was performed using 50% mobile phase A and 50% mobile phase B for 3 minutes. The flow rate was 0.30 mL/min, and the column temperature was maintained at 40°C. Targeted selected ion monitoring (t-SIM) data were acquired at a resolution of 35,000 with an AGC target of 5 x 10^4^, a maximum IT of 50 ms, and an isolation window of 1.0 m/z.

### Seahorse Analysis

Seahorse Mito Stress analysis was performed according to manufacturer’s instructions (Agilent). In brief, primary mouse gliomasphere cells were placed into an XF96e cell culture microplate at 50,000 cells/well on the day of the experiment in Seahorse XF DMEM Basal Medium supplemented with 10 mM glucose, 2 mM L-glutamine, and 1 mM sodium pyruvate and preincubated for at least 1 hour in a 37°C humidified CO2-free incubator. For αKG experiments, cells were pretreated with 10 mM αKG for at least 48 hours prior to the assay. To maintain conditions, the assay medium for these samples was also supplemented with 10 mM αKG. OCR was measured by a Seahorse XF96e Analyzer with the Mito Stress Test kit (Agilent). Mitochondrial function was interrogated by sequential injection of oligomycin (2 μM), FCCP (1 μM), and a rotenone/antimycin A mix (0.5 μM). Key mitochondrial parameters were calculated as follows: Basal respiration was derived from the basal OCR subtracted by post-rotenone/antimycin A OCR; ATP production was calculated as basal OCR subtracted by post-oligomycin OCR; and Maximal respiration was determined as the post-FCCP OCR subtracted by post-rotenone/antimycin A OCR.

### Single-cell RNA Sequencing Data Analysis

Single-cell RNA-seq data were obtained from GSE278456. Cells were annotated using same clustering and annotation schemes from the original study (*33*), including malignant cells, myeloid populations, stromal cells, and other immune cells. For SWELL1 stratification, patient-level SWELL1 expression was calculated, and patients were classified into high-and low-expression groups using the median as the cutoff. Single-cell data were aggregated into patient-level pseudobulk profiles, and differential expression was performed separately for each cell type at the patient level using DESeq2. P values were adjusted for multiple testing using the Benjamini– Hochberg procedure. Genes with FDR < 0.05 and |log2FC| > 1 were considered significant and subsequent GSEA analysis was performed using R “clusterProfiler” package(*81*), and terms with adjusted p value < 0.05 were considered significant.

### TCGA Database Analysis

TCGA database analyses were performed as previously described, with the following additions. All analyses utilized IDH-mutant lower-grade glioma (LGG) samples from the TCGA-LGG cohort (n = 427 IDH-mutant samples unless otherwise specified).

DNA methylation β-values (Illumina HumanMethylation450, level 3) and matched RNA-seq data [log2(TPM+1)] for TCGA-LGG samples (n = 530) were obtained from the UCSC Xena platform, and IDH-mutant samples (n = 427) were identified by unsupervised k-means clustering (k = 4). Per-sample mean β-values were computed over two CpG probe sets defined by Ceccarelli et al. (*4*): 1,308 IDH-mutant-specific probes, derived by excluding probes methylated in non-tumor brain (mean β ≥ 0.3) and selecting the top 5% most variable probes (1,308/25,978) within the IDH-mutant cohort, and 131 probes distinguishing G-CIMP-high from G-CIMP-low tumors (FDR < 10⁻¹⁵, |Δβ| > 0.27). Samples were stratified into *SWELL1*-High, *SWELL1*-Low, *LRRC8C*-High and *LRRC8C*-Low groups by median expression; group differences in mean β-values were assessed by Wilcoxon rank-sum test and continuous associations by Spearman rank correlation. *LRRC8C* promoter methylation was assessed using three CpG probes within the *LRRC8C* transcription start site CpG island (cg18312997, cg21541030, cg06353330). Analyses were performed in R using the IlluminaHumanMethylation450kanno.ilmn12.hg19 package.

Single-sample gene set enrichment analysis (ssGSEA) was performed with the GSVA R package using the HALLMARK_EPITHELIAL_MESENCHYMAL_TRANSITION and BENPORATH_SOX2_TARGETS gene sets retrieved from MSigDB. A G-CIMP-silenced gene set was curated from literature-established G-CIMP methylation targets (*MGMT, CDKN2A, RB1, MLH1, CDKN2B, RASSF1, DAPK1, CDH1*). Associations with *LRRC8C* expression were assessed by Spearman rank correlation and grade-adjusted partial Spearman correlation; group differences were assessed by Wilcoxon rank-sum test.

Tumor microenvironment cell composition was estimated from bulk RNA-seq data using the EPIC R package (*82*) in 394 IDH1-mutant LGG samples with available RNA-seq data. Purity-adjusted partial Spearman correlations between *LRRC8C* expression and estimated cell fractions were computed using TCGA tumor purity estimates. An 8-gene immunosuppression score was computed as the ranked expression of *TGFB2, IL10, IDO1, ARG1, CD274, IKZF2, HAVCR2*, and *CTLA4*, and a composite score was derived from a curated 27-gene immunosuppression panel. Multiple testing correction used the Benjamini-Hochberg FDR procedure.

For survival and disease-entity analyses, GDC-harmonized STAR-Counts RNA-seq data were obtained from the NCI Genomic Data Commons. Tumor purity was estimated by ABSOLUTE (*83*), and overall survival (OS), defined as time from diagnosis to death with living patients censored at last follow-up, was obtained from the TCGA Clinical Data Resource. For the genome-wide Cox proportional hazards screen, expression was rank-normalized [rank(TPM)/(n+1)], and each of 17,639 expressed genes (median TPM ≥ 1) was fit as OS ∼ expression + WHO grade + age at diagnosis + tumor purity. Gene-level significance was assessed by Wald test. For Kaplan-Meier analysis, patients were stratified by median *LRRC8C* expression and compared by log-rank test. For disease-entity analyses, tumors from TCGA-LGG were grouped according to available TCGA molecular annotations as IDH-mutant, 1p/19q-codeleted oligodendroglioma or IDH-mutant, 1p/19q-non-codeleted astrocytoma, approximating current molecular disease entities using retrospective TCGA annotations. IDH mutation status was determined from nonsynonymous IDH1 and IDH2 variants in GDC aliquot-ensemble masked somatic mutation calls; 1p/19q codeletion was determined from GDC masked copy number segments, with a chromosome arm called lost when more than 50% of arm probes resided in segments with log2 segment mean below -0.3 and codeletion requiring loss of both arms. Histologic grade used the available TCGA annotation. Only primary tumor samples were analyzed, with one aliquot per sample. Expression [log2(TPM + 1)] was compared across groups by Kruskal-Wallis test followed by pairwise two-sided Wilcoxon rank-sum tests with Benjamini-Hochberg correction.

### Quantitative Real-Time PCR

Total RNA was isolated using TRIzol reagent (Invitrogen). 0.5-2 μg of total RNA was used to generate the cDNA using the High-Capacity cDNA Reverse Transcription Kit (Applied Biosystems). The reaction was run in QuantStudio 6 Real-Time PCR Systems (Applied Biosystems) using 0.2 μL of cDNA in a 15 μL reaction. Primers targeting specific genes were provided in the Supplemental Material Table and used in combination with SYBR green master mix (Applied Biosystems) according to manufacturer’s instruction. The quantification was collected as CT value. Calibrations and normalizations were performed using the 2^-ΔΔCT method. Z-scores were calculated from 2^-ΔCT values. *Gapdh* or *ACTB* was used as the reference gene for mouse or human samples, respectively.

### Protein Immunoblotting

For histone 3 methylation and acetylation, histones from cells were extracted using a standard acid-extraction protocol (*84*). In brief, cell pellets were collected and resuspended in hypotonic lysis buffer (10 mM Tris, 1 mM KCl, 1.5 mM MgCl2, and 1 mM DTT, pH 8.0) containing protease inhibitors (Roche) at 4°C for 30 minutes followed by centrifugation at 10,000 xg for 10 minutes to remove cytosolic components. The nuclei were then resuspended in acid solution (0.4N HCl), and histones were extracted overnight at 4°C, followed by centrifugation at 16,000 xg for 10 minutes at 4°C. The supernatants were collected, and the histone pellets were precipitated with 35% (final concentration) trichloroacetic acid on ice. After washing with ice-cold acetone and drying at room temperature, the histone pellet was dissolved in H2O and prepared for immunoblotting.

Whole cell lysates were prepared in RIPA buffer with protease inhibitor cocktails (Roche). Lysates in RIPA buffer were cleared of debris by centrifugation at 12,000 xg for 10 minutes at 4°C. The concentration of whole cell lysates or isolated histones was determined by BCA assay kit (Thermo Fisher Scientific). Lysates and histones were then collected and made into same concentrations with water and Blot 4x LDS sample buffer (Invitrogen) before boiling at 75°C for 10 minutes prior to SDS-PAGE electrophoresis. Novex 4-20% gradient gel (Invitrogen) was used for electrophoretic separation according to manufacturer’s instruction. For subsequent antibody detection, nitrocellulose membranes were used for protein transfer. Primary antibodies were made according to manufacturers’ instruction and incubated with nitrocellulose membranes overnight at 4°C. Following primary antibody incubation, samples were incubated with HRP-conjugated anti-mouse, anti-rabbit, or anti-rat secondary antibody (Proteintech) for 1-2 hours at room temperature for the visualization.

### Statistics

All data were analyzed using GraphPad Prism or RStudio. Data are presented as mean with standard error of the mean (SEM) as indicated in figure legends. Survival curves are presented as Kaplan-Meier plots and compared by log-rank (Mantel-Cox) test. Group comparisons were performed using unpaired two-tailed Student’s t-test, Welch’s t-test, one-way ANOVA, or two-way ANOVA followed by Sidak’s multiple comparisons test, as indicated in figure legends. Non-normal or ordinal data were analyzed by Wilcoxon rank-sum test or Kruskal-Wallis test with pairwise Wilcoxon rank-sum tests and Benjamini-Hochberg correction. Correlations were assessed by Spearman rank correlation, with partial Spearman correlation (ppcor) used for covariate adjustment. Differential expression and Cox regression gene-level significance were assessed by Wald test with Benjamini-Hochberg correction where applicable. Linear and non-linear regressions were performed in GraphPad Prism. A p value less than 0.05 was considered statistically significant. Sample sizes are indicated in figures and legends.

