## Supplementary Figures for "SWELL1 channel-mediated D-2HG export promotes immune evasion and metabolic fitness in IDH-mutant glioma"

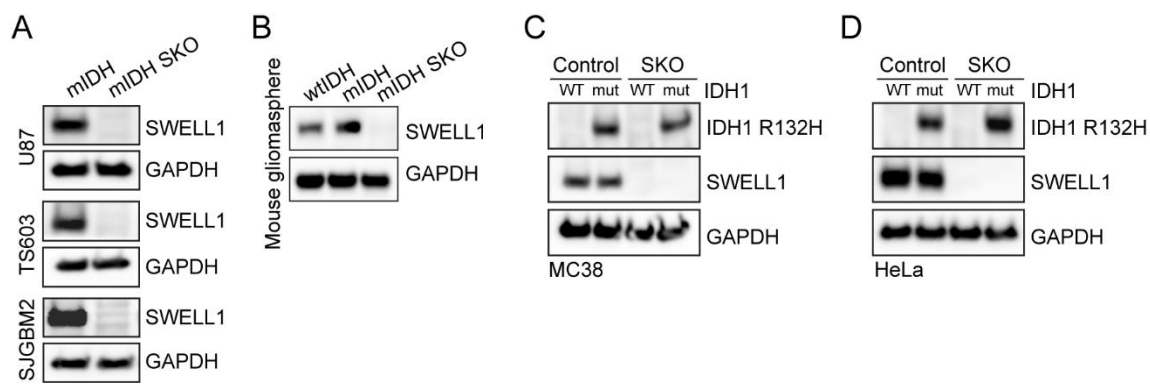

**Figure S1. Generation of mIDH SWELL1 KO cells.**

**A–D,** Representative immunoblotting of mIDH-KI U87, TS603, mIDH-transduced SJGBM2 (A), mouse gliomasphere cells (B), mIDH-transduced MC38 (C), or mIDH-transduced HeLa cells (D) for the validation of SWELL1 deletion or mIDH expression (n = 1). mut-IDH1 in (C–D) refers to mIDH in Fig 1L. TS603 in (A) is shared with Fig. 5C.

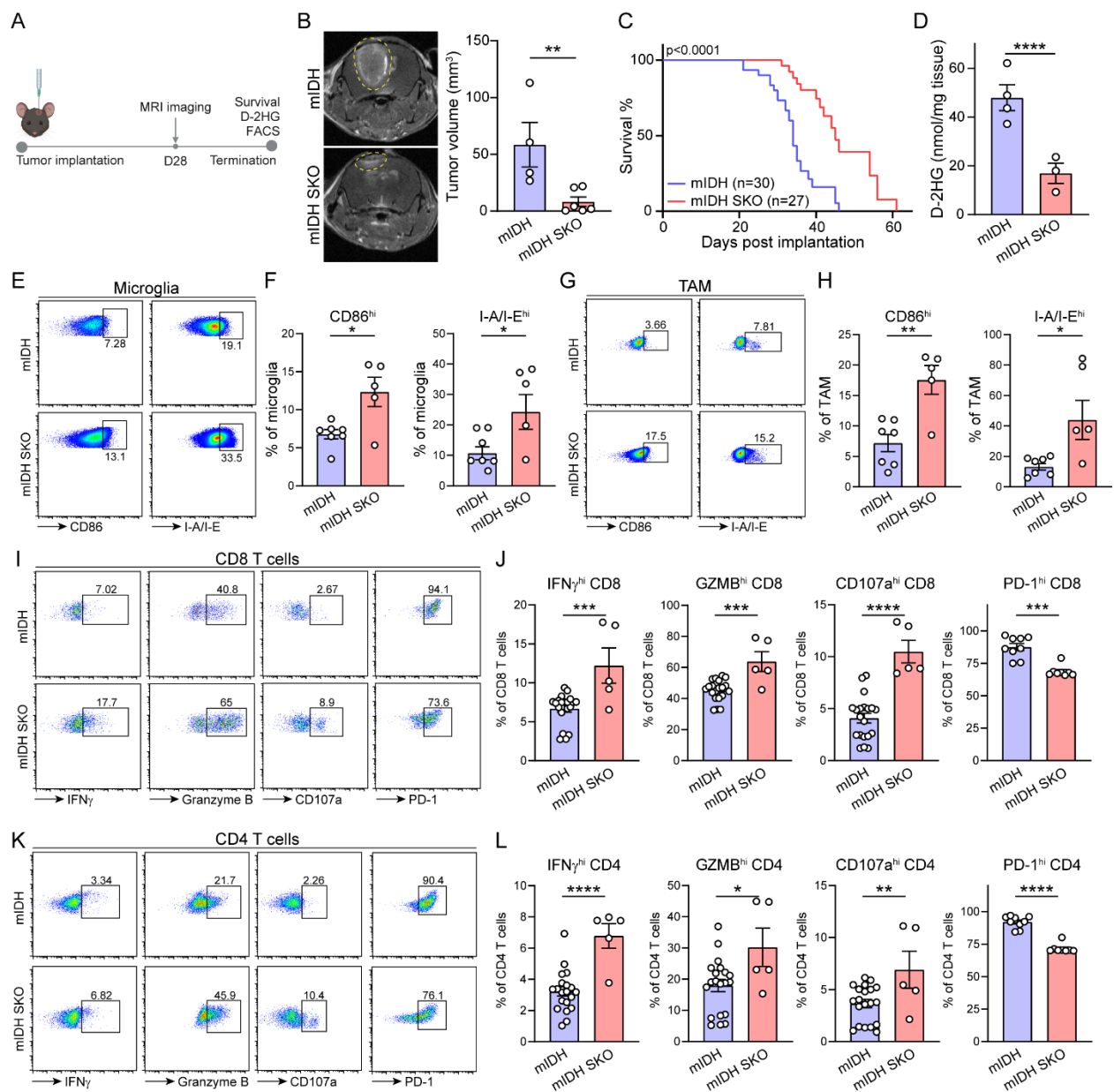

**Figure S2. SWELL1 deletion attenuates D-2HG-driven immunosuppression in syngeneic mouse IDH-mutant gliomas.**

**A**, Serial sections of the representative MRI images shown in **Fig. 2B**.

**B–C**, FACS gating strategy for tumoral CD45<sup>+</sup> immune cells (**B**) and myeloid populations (**C**).

**D**, FACS quantification of tumor-associated neutrophils (TAN), microglia, monocytic myeloid-derived suppressor cells (m-MDSC), and tumor-associated macrophages (TAM) populations in mouse gliomasphere tumors (n = 8 to 10).

**E–F**, FACS quantification of intracellular arginase 1 and surface PD-L1 expression in tumoral microglia (**E**) or TAMs (**F**) (n = 8 to 10).

**G**, FACS gating strategy for tumoral T cell populations following gating in **(B)**.

**H**, FACS quantification of CD8<sup>+</sup> T, regulatory T (Tregs), and not-Treg CD4<sup>+</sup> T cell populations in mouse gliomasphere tumors (n = 6 to 9).

**I–J**, FACS quantification of intracellular IL-2 and TNFα expression within tumoral CD8<sup>+</sup> T (**I**) or CD4<sup>+</sup> T (**J**) cells (n = 8 to 9).

**K**, Variance-stabilizing transformation (VST) normalized expression (log<sub>2</sub> scale) of raw DESeq2-normalized counts of curated immunogenicity genes across three functional modules in the RNA-seq results (n = 3).

Data are reported as mean ± SEM. Unpaired t-test for **D**, **E**, **F**, **H**, **I**, **J**. Wald test with Benjamini-Hochberg correction for **K**. \*p < 0.05, \*\*p < 0.01, \*\*\*p < 0.001, and \*\*\*\*p < 0.0001.

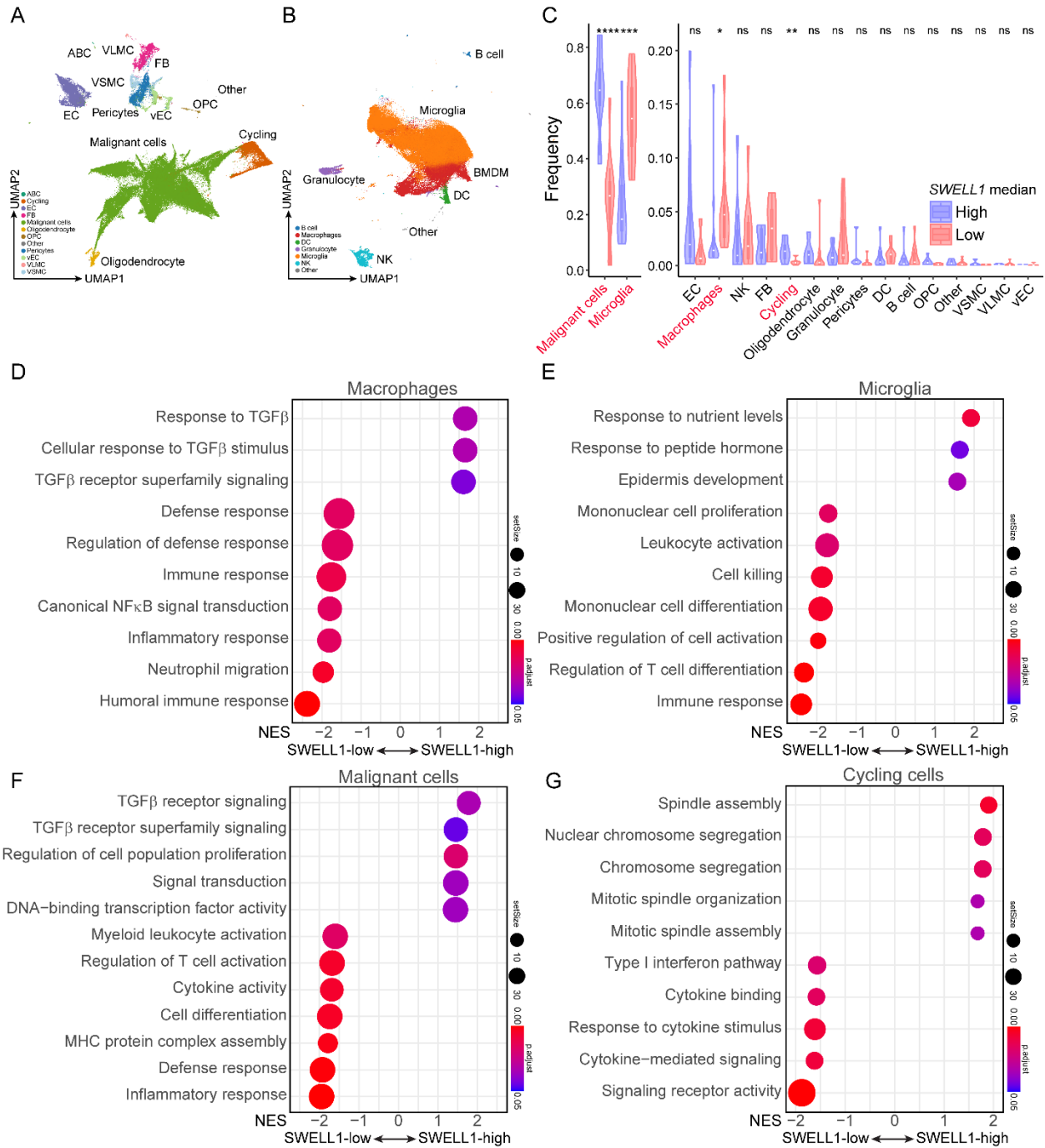

**Figure S3. Low SWELL1 expression correlates with an immunoactive TME in human IDH-mutant gliomas.**

**A–B**, UMAP projections of integrated dataset (GSE278456) from pooled samples for CD45– non-immune cells (**A**) and CD45+, CD3– cells (**B**). Total of 123,647 cells within 12 IDH-mutant tumors.

**C**, Relative cell population frequencies identified within mIDH gliomas, calculated per patient. SWELL1-high and SWELL1-low groups were defined by the median SWELL1 expression in malignant cells.

**D–G**, Selected gene set enrichment analysis (GSEA) for macrophages (**D**), microglia (**E**), malignant cells (**F**), or cycling cells (**G**) within mIDH gliomas. SWELL1-high/low groups were defined as in **C**.

Violin plots show the distribution of per-patient frequencies. Wilcoxon rank-sum test was used for **C**. ns, $p > 0.05$, $*p < 0.05$, $**p < 0.01$, and $***p < 0.001$.

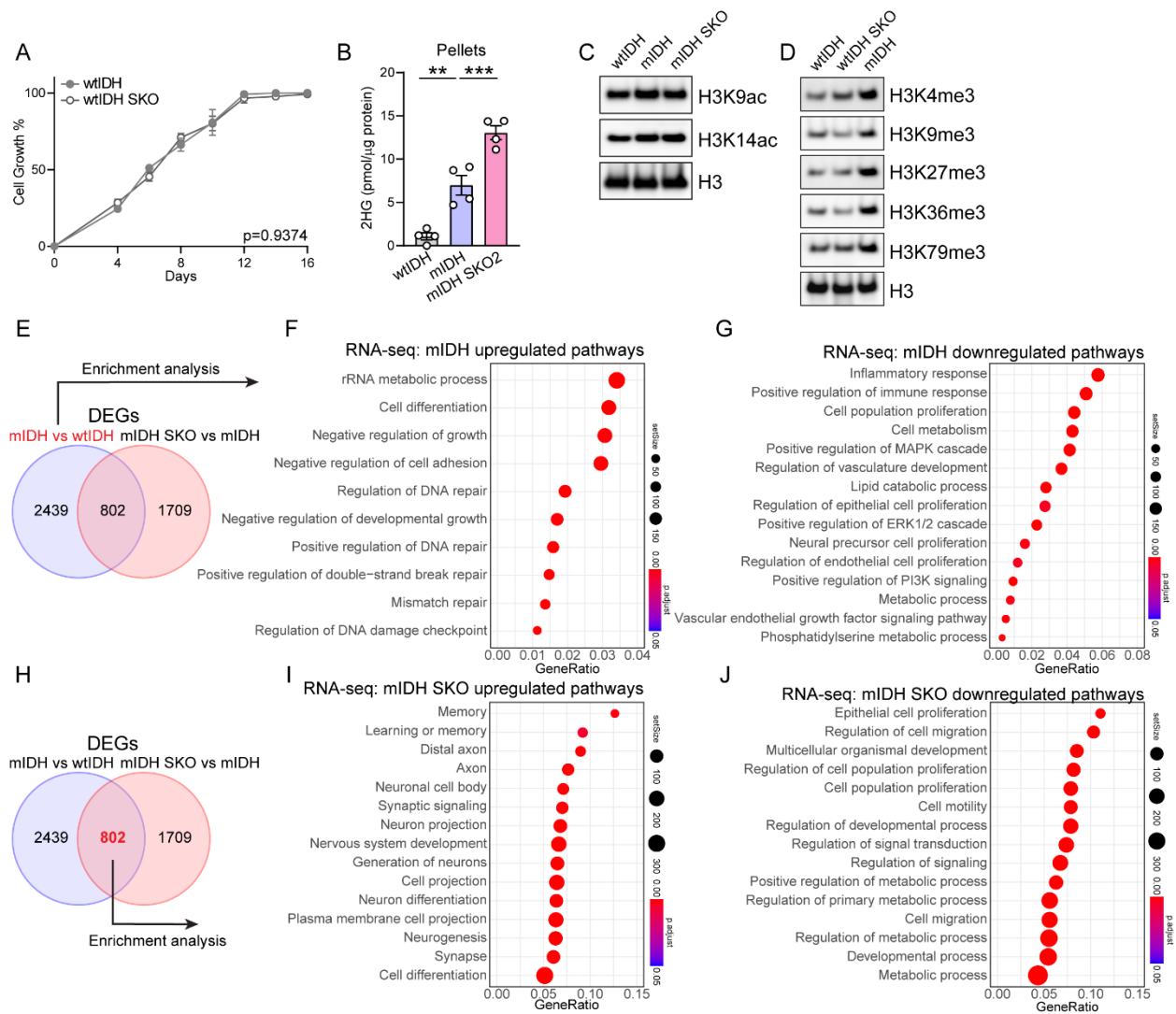

**Figure S4. VRAC-mediated D-2HG efflux shapes D-2HG-driven transcriptional changes in mIDH glioma cells.**

**A**, Colony formation assay for mouse gliomasphere cells. Data normalized to the maximum absorbance within the comparison (n = 3). wtIDH is shared with **Fig. 3B**.

**B**, 2HG concentration normalized to total protein levels in mouse gliomasphere cell pellets (n = 4). wtIDH and mIDH are shared with **Fig. 3C**.

**C**, Immunoblotting of isolated histones from mouse gliomasphere cells for H3 acetylations (n = 1). H3 is shared with **Fig. 3D**.

**D**, Immunoblotting of isolated histones from mouse gliomasphere cells for H3 trimethylations (n = 3).

**E**, Venn graph comparison of differential expressed gene (DEGs) numbers from RNA-seq analysis with comparisons of mIDH vs wtIDH and mIDH SKO vs mIDH (n = 3). Highlighted mIDH vs wtIDH comparison was used for analysis in (**F**) and (**G**).

**F–G**, Selected gene ontology analysis for upregulated (**F**) or downregulated (**G**) genes from RNA-seq analysis (n = 3).

**H**, Venn graph comparison of differential expressed gene (DEGs) numbers from RNA-seq analysis with comparisons of mIDH vs wtIDH and mIDH SKO vs mIDH (n = 3). Highlighted DEGs were used for analysis in (**I**) and (**J**).

**I–J**, Selected gene ontology analysis for upregulated (**I**) or downregulated (**J**) genes from RNA-seq analysis (n = 3).

Data are reported as mean ± SEM. Two-way ANOVA with Sidak's test for **A**. One-way ANOVA with Sidak's test for **B**. \*\*p < 0.01 and \*\*\*p < 0.001.

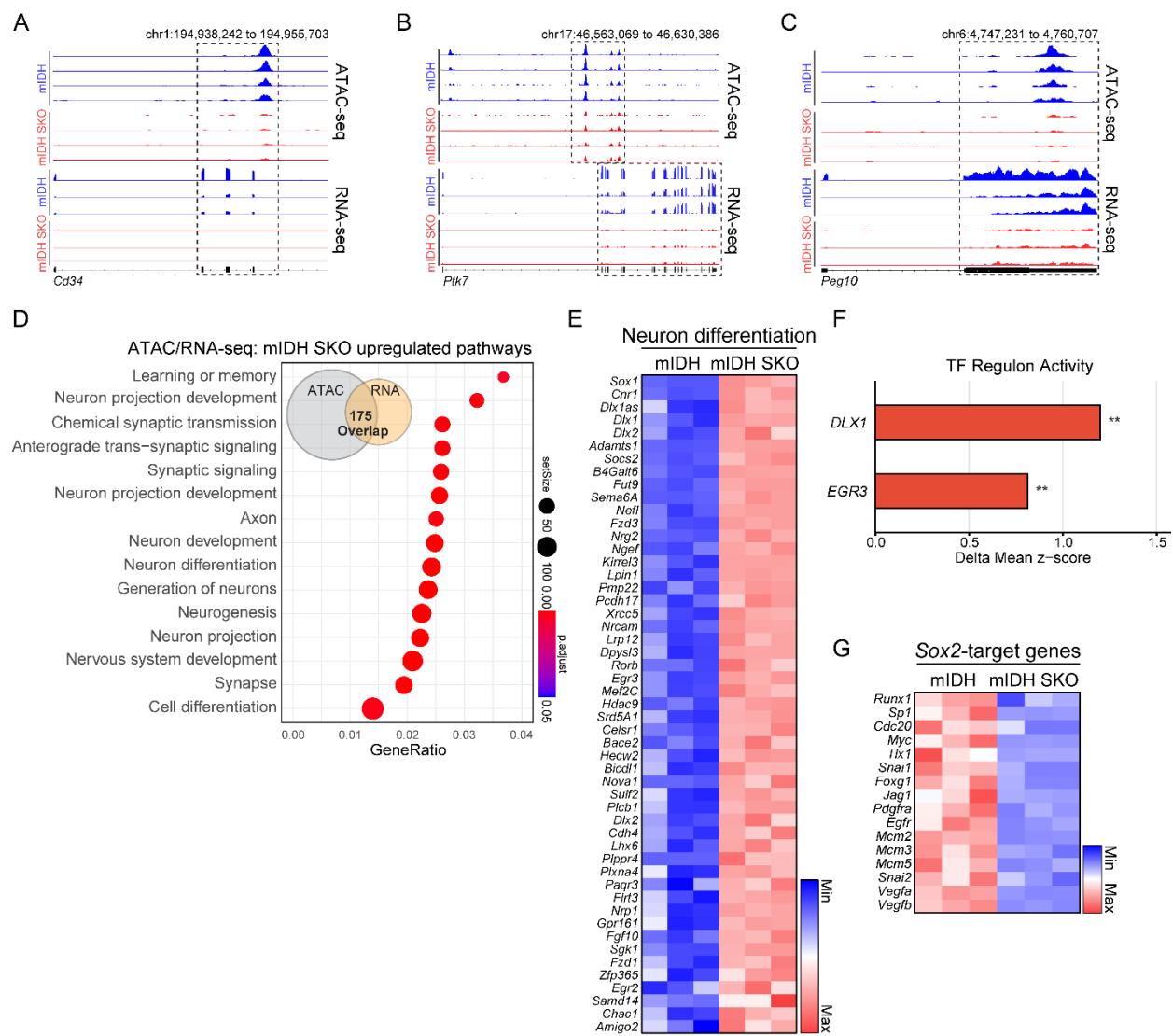

**Figure S5. VRAC-mediated D-2HG efflux shapes D-2HG-driven epigenetic remodeling in mIDH glioma cells.**

**A–C**, IGV profiles of ATAC-seq and RNA-seq signals for *Cd34* (**A**), *ptk7* (**B**), or *Peg10* (**C**) gene in mouse gliomasphere cells (ATAC-seq: $n = 4$; RNA-seq: $n = 3$). Loci are oriented such that the TSS is shown on the left.

**D**, Selected gene ontology analysis from integrated ATAC-seq and RNA-seq analysis for upregulated pathways. 175 upregulated genes shared between ATAC-seq and RNA-seq were used for the analysis (ATAC-seq: $n = 4$; RNA-seq: $n = 3$).

**E**, Heatmaps of cell differentiation gene expressions within mouse gliomasphere RNA-seq data. Data presented in z-scores calculated from FPKM ($n = 3$).

**F**, Transcription factor (TF) regulon activity change. Regulon activity was scored as the mean z-score of curated target genes for each TF across all six samples. Bars represent the difference in mean regulon z-score between groups (delta mean z-score).

**G**, Heatmaps of Sox2-target gene expressions within mouse gliomasphere RNA-seq data. Data presented in z-scores calculated from FPKM (n = 3).

Welch's t-test for **F**. \*\*p < 0.01

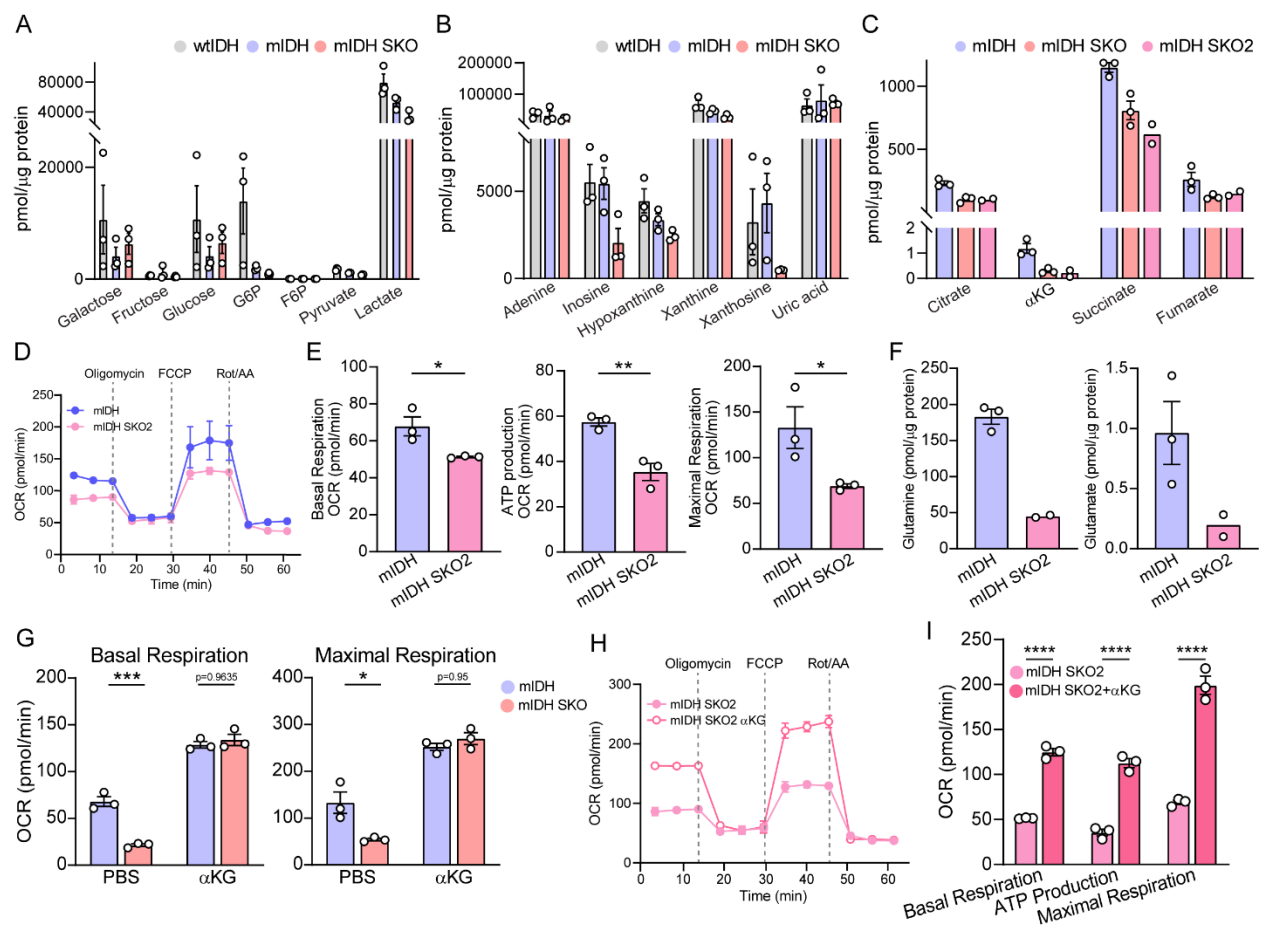

**Figure S6. VRAC-mediated D-2HG release alleviates mitochondrial metabolic stress in mIDH cells.**

**A–B**, Glycolysis (**A**) or purine catabolism (**B**) metabolite concentrations normalized to total protein levels in mouse gliomasphere cell pellets (n = 3).

**C**, TCA cycle metabolite concentrations normalized to total protein levels in mouse gliomasphere cell pellets (mIDH and mIDH SKO: n = 3; mIDH SKO2: n = 2). mIDH and mIDH SKO are shared with **Fig. 4A**.

**D–E**, Seahorse analysis of cellular oxygen consumption rate (OCR) kinetics. Representative OCR trace (**D**) and quantifications (**E**) measured in mouse gliomasphere cells (n = 3). mIDH is shared with **Fig. 4B–C**. mIDH SKO2 is shared with **Fig. S6I**.

**F**, Glutamine and glutamate concentrations normalized to total protein levels in mouse gliomasphere cell pellets (mIDH: n = 3; mIDH SKO2: n = 2). mIDH is shared with **Fig. 4D**.

**G**, Seahorse analysis of cellular oxygen consumption rate (OCR) quantifications measured in mouse gliomasphere cells supplemented with 10 mM αKG (n = 3). Data corresponds to **Fig. 4F–G**. Untreated mIDH and mIDH SKO are shared with **Fig. 4B–C**.

**H–I**, Seahorse analysis of cellular oxygen consumption rate (OCR) kinetics. Representative OCR trace (**H**) and quantifications (**I**) measured in mouse gliomasphere cells supplemented with 10 mM αKG (n = 3). untreated mIDH SKO2 is shared with **Fig. S6D–E**.

Data are reported as mean ± SEM. Unpaired t-test for **E**. Two-way ANOVA with Sidak's test for **G, I**. \*p < 0.05, \*\*p < 0.01, and \*\*\*p < 0.001.

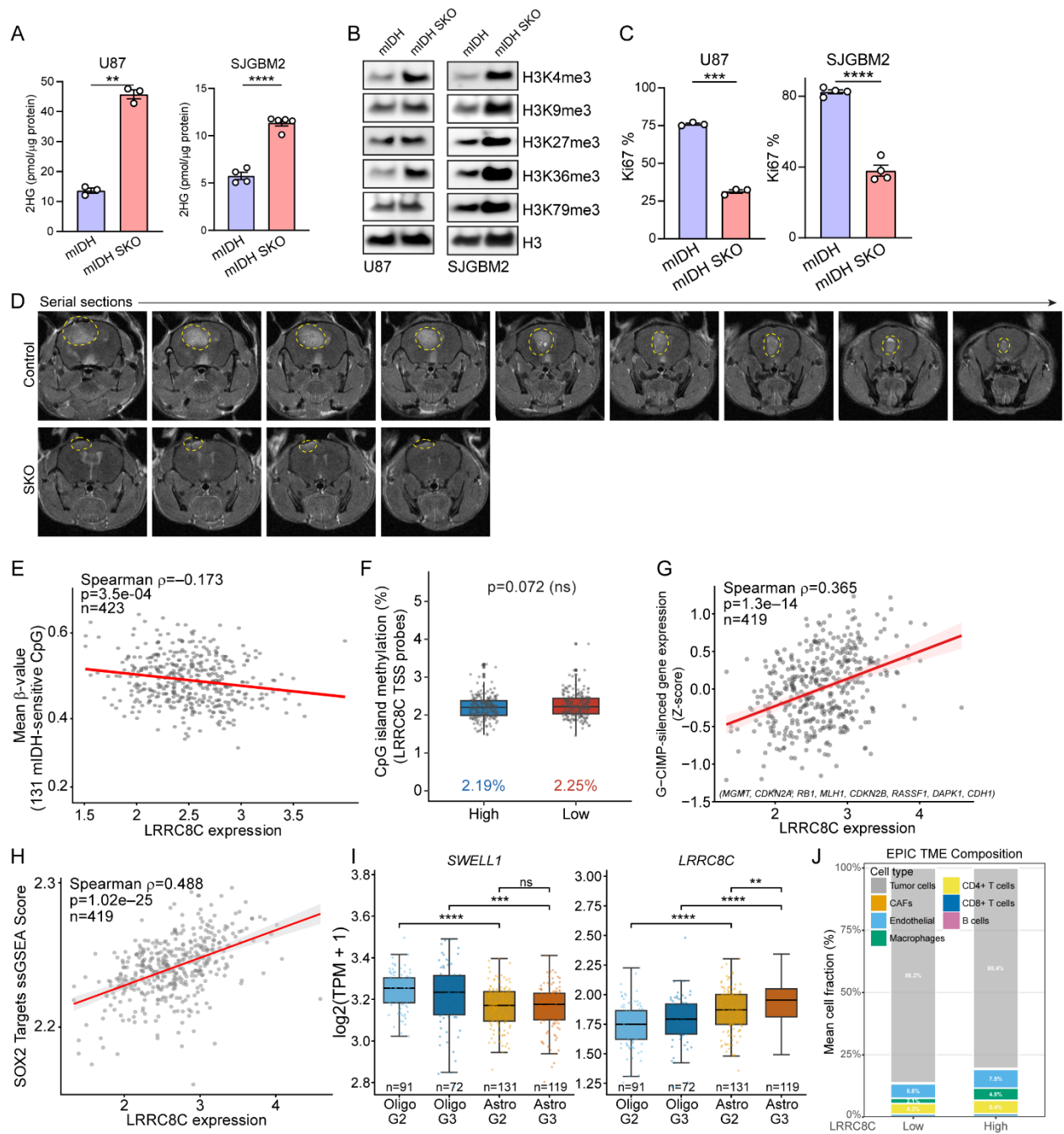

**Figure S7. SWELL1 deletion suppresses cell growth in human mIDH glioma cells.**

**A**, 2HG concentration normalized to total protein levels in U87 (left) or SJGBM2 (right) cell pellets (n = 3 to 5). U87 mIDH is shared with Fig. 1A.

**B**, Immunoblotting of isolated histones from U87 (left) or SJGBM2 (right) cells for H3 trimethylations (n = 2).

**C**, FACS quantification of Ki67 positivity of U87 (left) or SJGBM2 (right) cells (n = 3 to 4).

**D**, Serial sections of the representative MRI images shown in **Fig. 5I**.

**E**, Scatter plot showing the correlation between *LRRC8C* expression ($\text{Log}_2(\text{TPM}+1)$) and mean DNA methylation beta-value at 131 mIDH-sensitive CpG sites in IDH-mutant LGG samples from TCGA-LGG database.

**F**, Box plot comparing CpG island methylation levels at *LRRC8C* TSS probes between *LRRC8C*-high and -low IDH-mutant LGG tumors.

**G**, Scatter plot showing the correlation between *LRRC8C* expression ($\text{Log}_2(\text{TPM}+1)$) and the expression of a curated G-CIMP-silenced gene set in IDH-mutant LGG samples from TCGA-LGG database.

**H**, Scatter plot showing the correlation between *LRRC8C* expression ($\text{Log}_2(\text{TPM}+1)$) and SOX2 target gene activity (ssGSEA score) in IDH-mutant LGG samples from TCGA-LGG database.

**I**, Box plots with individual data points showing *LRRC8A* (SWELL1, left) and *LRRC8C* (right) expression ($\text{log}_2(\text{TPM}+1)$) across the four WHO CNS5 IDH-mutant glioma entities in IDH-mutant LGG samples from the TCGA-LGG database: oligodendroglioma, IDH-mutant and 1p/19q-codeleted, grade 2 (n = 91) and grade 3 (n = 72), and astrocytoma, IDH-mutant and non-codeleted, grade 2 (n = 131) and grade 3 (n = 119). Boxes show the median and interquartile range; whiskers extend to 1.5x the interquartile range.

**J**, Stacked bar chart of EPIC-estimated tumor microenvironment (TME) cell type composition in *LRRC8C*-low versus *LRRC8C*-high IDH-mutant LGG tumors.

Data are reported as mean ± SEM. Unpaired t-test for **A**, **C**. Wilcoxon rank-sum test for **F**. Spearman rank correlation for **E**, **G**, **H**. Kruskal-Wallis test with pairwise two-sided Wilcoxon rank-sum tests, Benjamini-Hochberg correction, for **I**. \*\*p < 0.01, \*\*\*p < 0.001, and \*\*\*\*p < 0.0001. ns, not significant.

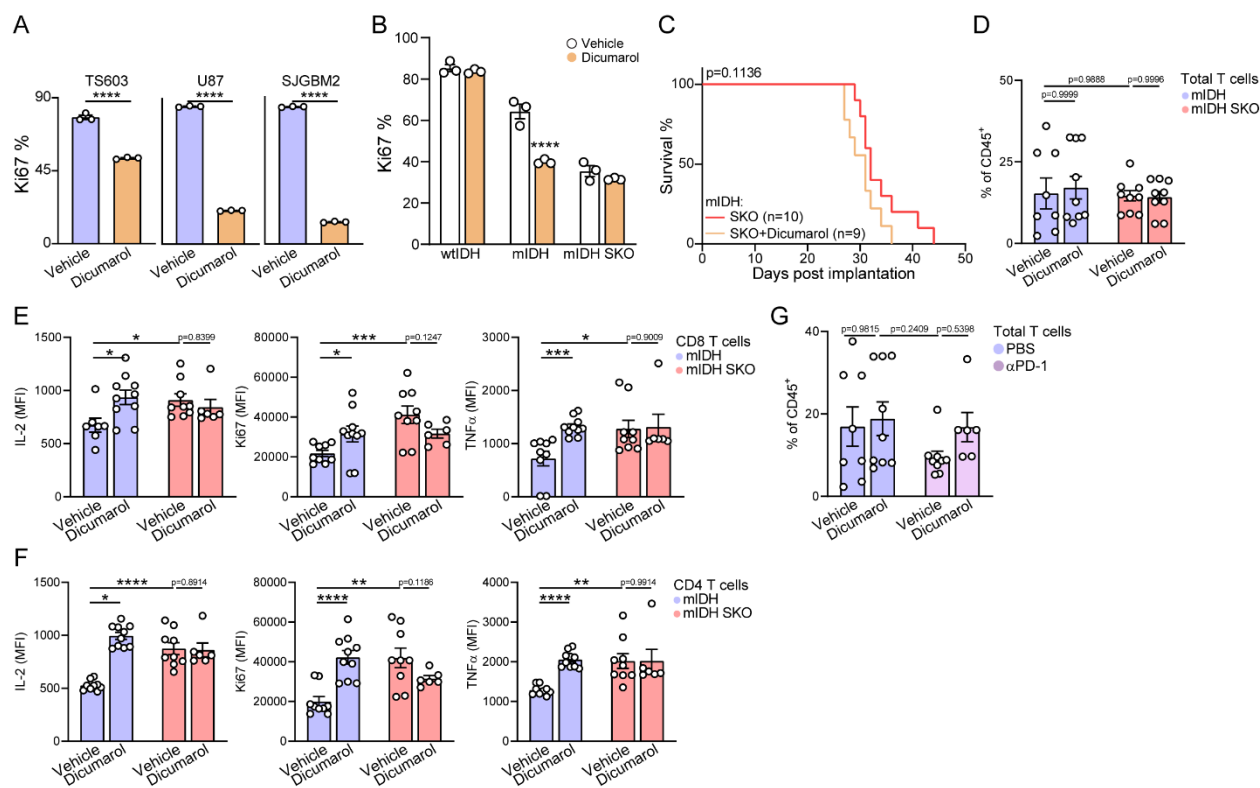

**Figure S8. Pharmacological VRAC inhibition enhances antitumor immunity in syngeneic mouse IDH-mutant gliomas.**

**A**, FACS quantification of Ki67 positivity of TS603 (left), U87 (mid), or SJGBM2 (right) cells treated with dicumarol (n = 3).

**B**, FACS quantification of Ki67 positivity of mouse gliomasphere cells treated with dicumarol (n = 3). Vehicle-treated samples were used in **Fig. 3A**.

**C**, Survival of mice implanted with $7 \times 10^4$ mouse gliomasphere cells treated with dicumarol.

**D**, FACS quantification of T cells in mouse gliomasphere tumors treated with dicumarol (n = 8 to 10). mIDH vehicle and dicumarol are shared with (**G**).

**E–F**, FACS quantification of intracellular IL-2, Ki67, and TNFα expression within tumoral CD8+ T (**E**) or CD4+ T (**F**) cells with dicumarol treatments (n = 6 to 7).

**G**, FACS quantification of T cells in mouse gliomasphere tumors treated with dicumarol and anti-PD1 antibody (n = 6 to 10). αPD-1, anti-PD1 antibody. PBS vehicle and dicumarol are shared with (**D**).

Data are reported as mean ± SEM. Unpaired t-test for **A**, **B**. Mantel-Cox test for **C**. Two-way ANOVA with Sidak's test for **D**, **E**, **F**, **G**. \*p < 0.05, \*\*p < 0.01, \*\*\*p < 0.001, and \*\*\*\*p < 0.0001.
